# High-PepBinder: A pLM-Guided Latent Diffusion Framework for Affinity-Aware Target-Specific Peptide Design

**DOI:** 10.64898/2026.01.12.698988

**Authors:** Qingyi Mao, Silong Zhai, Sen Cao, Renjie Zhu, Wen Xu, Chengyun Zhang, Ning Zhu, Jingjing Guo, Hongliang Duan

## Abstract

Peptides, as therapeutic molecules, offer unique advantages in targeting complex protein surfaces, yet their rational design remains limited by the vastness of the sequence space and the constraints of traditional approaches. Here, we propose High-PepBinder, a sequence-only conditional diffusion framework for target-specific peptide generation. Guided by the target protein sequence, High-PepBinder adopts a dual encoder architecture that integrates protein language models (pLMs) with the diffusion model. This approach cascades the peptide generation model with an affinity classifier and enables the generation process to capture affinity-related features of the peptides through lightweight joint optimization. Due to the scarcity of protein-peptide affinity data, we constructed PepPBA, to our knowledge the most comprehensive dataset to date, and established a structure- and physics-based screening pipeline to prioritize top candidates. Results show that High-PepBinder demonstrates competitive performance across multiple peptide generation and affinity-related tasks. For representative targets, including KEAP1, XIAP, and EGFR, the generated peptides preserve key binding geometries and interface patterns of reference peptides in predicted complexes, while maintaining sequence diversity and favorable predicted properties. Overall, High-PepBinder contributes toward a general and sequence-only strategy for peptide design, offering a computational framework for expanding peptide discovery against challenging targets.

## 1. Introduction

Many pathogenic proteins lack stable small-molecule binding pockets, and although antibodies offer high specificity, their large size prevents them from entering cells, limiting their applicability to intracellular targets^1–5^. In contrast, peptides combine a moderate molecular size, favorable biophysical properties, and the ability to engage large protein surfaces^6–9^. Their flexible conformational space enables them to recognize complex topologies and efficiently target traditionally “undruggable” proteins, including intrinsically disordered proteins and protein-protein interactions (PPIs) ^10^. However, a major challenge in designing target-specific peptides is the need to explore an enormous chemical space. Traditional peptide discovery relies on phage or yeast display to screen mutation libraries, a process that is time-consuming, costly, and labor-intensive^11,12^. Another class of strategies, such as alanine scanning, is more efficient but limited by its dependence on prior structural information and its restricted ability to explore sequence diversity^13^. These limitations further underscore the urgent need for more effective peptide engineering approaches.

Recently, a large number of data-driven deep learning models have emerged to reduce the cost and time required for peptide design. These models can be categorized into two types: predictive models and generative models^14^. Predictive models aim to evaluate protein-peptide binding strength or interaction likelihood^15,16^. Although effective, they typically rely on existing sequence databases or experimental samples and lack the ability to explore novel sequence space. As a result, they often require additional heuristic searches or mutation scans, which limits their efficiency. In contrast, generative models seek to learn latent distributions from data to expand the accessible sequence space^17–19^. However, their generation process is often decoupled from functional objectives and lacks explicit mechanisms for functional optimization. Generative models can be further divided into sequence-based methods (e.g., PepMLM) and structure-sequence co-design methods (e.g., RFdiffusion + ProteinMPNN)^20–22^. In many practical applications, obtaining high-quality target structures is challenging, and target proteins may exhibit dynamic conformations. Although structure-sequence co-design methods can leverage structural prior information, they typically rely on the 3D structure of binding sites, limiting their applicability in the absence of structural information or when the target is highly dynamic. This has led to the adoption of sequence-based methods, which offer broader applicability and scalability. However, most sequence-based generative models fail to effectively incorporate affinity-related signals into the generation process in the absence of reliable functional supervision. Consequently, there remains a need for a structure-independent algorithm that can introduce affinity-aware constraints during generation, enabling the design of target-conditioned peptides.

The development of protein language models (pLMs) offers a new opportunity to address this challenge^23–26^. These models can capture both sequence and structural signals within a unified representation space, thereby providing a natural representation space that is well suited for latent diffusion-based generative modeling^27^. In this context, we introduce High-PepBinder, a sequence-only peptide design framework tailored for target-specific applications (Figure 1b). High-PepBinder integrates pLM representations with a diffusion-based generative model to achieve target-conditioned peptide sequence generation and affinity-aware optimization within a unified end-to-end architecture. Affinity awareness is introduced through two complementary mechanisms: target protein conditional modeling and a predictive module that acts as a functional constraint jointly guiding the generation process. Through conditional encoding, target protein sequence representations are injected into the diffusion denoising procedure, enabling the model to learn target-specific binding preferences and interaction priors without relying on explicit 3D structural inputs, thereby supporting peptide generation tailored to specific targets. To enhance modeling across diverse targets, High-PepBinder adaptively optimizes peptide sequence representations during training, allowing generic pLM representations to better align with peptide generation and to effectively exploit target-conditioned context. Furthermore, to more tightly couple conditional generation with functional objectives, High-PepBinder introduces a lightweight joint fine-tuning stage and applies consistency regularization to improve the stability of the predictive module under diffusion noise perturbations, thereby strengthening the consistency and generalization of the generation-screening workflow^28^.

Because of the scarcity of protein-peptide affinity datasets, we constructed PepPBA, a highly comprehensive protein-peptide affinity dataset that substantially surpasses existing resources in both size and target coverage^29,30^. We also developed a complementary posterior validation pipeline that evaluates the structural plausibility and energetic stability of predicted complexes by integrating AlphaFold3-based scoring, Rosetta binding energy calculations, and molecular dynamics (MD) simulations to support post hoc in silico evaluation of structural plausibility and energetic stability^31–38^. We validated the robustness of High-PepBinder across multiple benchmarks and three high-value therapeutic targets, EGFR, KEAP1 and XIAP. High-PepBinder demonstrates competitive or improved performance relative to RFdiffusion and PepMLM in peptide generation quality and high-affinity enrichment across the evaluated benchmarks. Moreover, its affinity classifier achieves competitive accuracy under cold-start splits, demonstrating strong generalization and reliable screening performance. Overall, High-PepBinder provides a unified framework for design target-specific peptide candidates, without requiring explicit target 3D structures as model inputs, offering a new avenue for drug discovery against traditionally undruggable targets.

**Figure 1.**
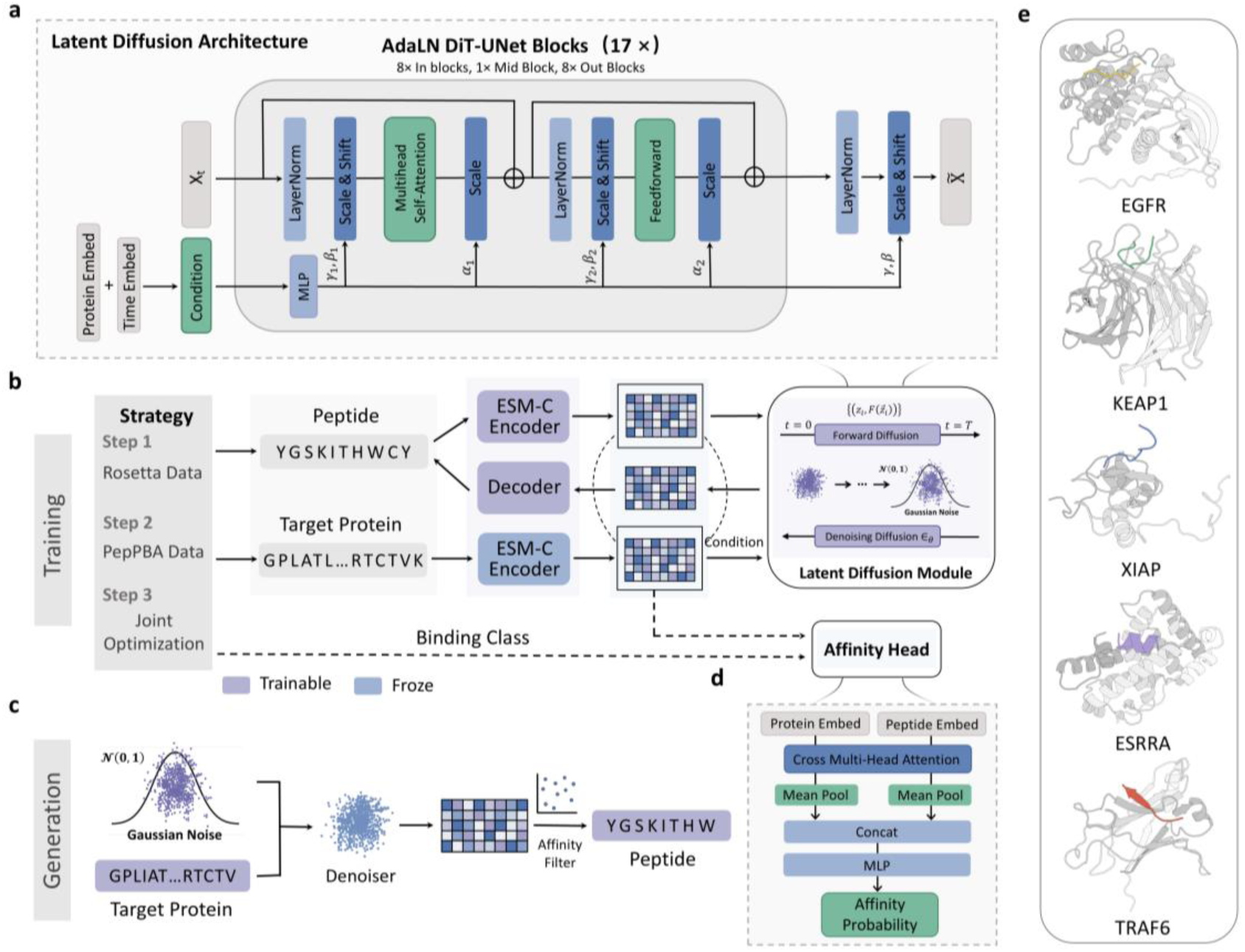
Overview of High-PepBinder: an integrated generation-prediction framework for target-specific peptide design. (a) Latent diffusion module. The model adopts a Transformer-UNet architecture consisting of 17 blocks, with 8 encoder layers, 1 bottleneck layer, and 8 decoder layers, and incorporates cross-layer skip connections for multilevel feature fusion. Protein sequence embeddings and diffusion timesteps are combined to form the conditional vector, which guides the network to generate high-dimensional latent representations of peptides. (b) Training pipeline. The training process comprises three stages: weakly supervised pretraining on Rosetta-scored pseudo protein-peptide pairs, finetuning on a protein-peptide affinity dataset, and a lightweight joint optimization fine-tuning stage to further enhance protein-conditioned peptide generation. (c) Inference pipeline. Target-specific peptide sequences are generated based on the input target information. (d) Lightweight affinity classifier. Architecture of the dual-encoder, cross-attention-based affinity prediction module.

## 2. Methods

### 2.1 High-PepBinder Framework

Figure 1b and 1c illustrate the architecture of High-PepBinder and the information flow during model training and generation. High-PepBinder is a two-stage cascade model composed of a generator and an affinity discriminator, each contributing complementary affinity-related information to enhance the learning process. The generator integrates three core components: an Encoder Block, a Latent Diffusion Block, and a Decoder Block (Figure 1b). High-PepBinder adopts a dual-encoder architecture based on the pretrained protein language model ESM Cambrian (ESM-C), in which the target sequence and peptide sequences are encoded separately^39^. Owing to differences in sequence length and statistical properties between proteins and peptides, we unfreeze the peptide-side encoder during training to enhance its adaptability to peptide-specific features and target-dependent affinity patterns. The Latent Diffusion Block operates in the learned peptide latent space and is implemented using a modified diffusion transformer architecture^40^ (Figure 1a). The target protein embedding is incorporated as conditional information and combined with the diffusion timestep to guide peptide generation. The resulting peptide latent representations are decoded into peptide sequences by the Decoder Block, which follows the MLP-based decoding architecture used in ESM-family models^26^. Finally, a lightweight cross-attention-based affinity classifier is applied to filter the generated sequences (Figure 1d).

The generative model is trained using a multi-stage strategy. First, the model is pretrained on large-scale weakly supervised pseudo protein-peptide pairs dataset to learn transferable interface priors. It is then fine-tuned on experimentally validated protein-peptide interaction data with affinity annotations, allowing the model to transition from weakly supervised interface representations to a functional space aligned with true affinity signals. Finally, we introduce a lightweight joint optimization fine-tuning stage to gently incorporate weakly supervised affinity signals into the generation process. Building on this, we further introduce a noise-consistent regularization auxiliary loss based on diffusion latent variables, which enhances the robustness and generalization of the classifier^41^.

#### 2.1.1 Protein-Peptide Encoders

To fully leverage the prior structural-functional representation capabilities of pLMs, we use the ESM-C as the foundational encoder for both protein and peptide sequences^39^. ESM-C is a parallel branch of the ESM-3 framework built on a multi-layer Transformer architecture and pretrained on large-scale protein sequence data. It is designed to learn evolutionary-scale protein representations across diverse domains of life, capturing both structural constraints and functional semantics at the sequence level. Compared with ESM2, ESM-C features increased model capacity and a substantially larger training corpus, enabling it to generate biologically richer embeddings and achieve superior performance in structure prediction and function inference tasks. The ESM-C encoder can be formally represented as a mapping function:

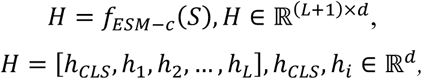

where 𝑑 = 1152 denotes the dimensionality of the hidden representations, ℎ_𝑖_ represents the residue-level contextual embedding, and ℎ_CLS_ denotes the hidden vector of the [CLS]token.

In this work, we employ two ESM-C encoders to process protein and peptide sequences separately:

- Protein channel: the ESM-C parameters are frozen to preserve evolutionary priors and prevent overfitting or degradation.
- Peptide channel: the ESM-C parameters are finetuned to adapt the model to the diffusion-based reconstruction task and enhance the generative quality of peptide representations.

For protein sequences, we construct a global conditional vector by concatenating the [CLS] token embedding with a sequence-level representation obtained via mean pooling over all residue embeddings:

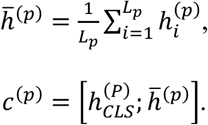

This global vector is then used to condition the diffusion module. For peptide sequences, we retain the token-level embeddings as the initial latent inputs to the diffusion network.

#### 2.1.2 Conditional Latent Diffusion

##### DiT-UNet Denoising Architecture

Unlike the standard DiT model, which adopts a unidirectional, layer-by-layer stacked architecture, High-PepBinder employs a DiT-UNet architecture that extends the diffusion Transformer into a symmetric encoder-bottleneck-decoder pathway and introduces cross-layer skip connections to enhance multilevel semantic representations and generative consistency (Figure 1a)^40^. The motivation behind this design is that, although peptide sequences lack explicit spatial resolution, their structure-function dependencies are hierarchical (primary-sequence motifs, local structural patterns, long-range interactions and so on). Thus, a UNet-style hierarchical fusion mechanism helps capture multi-scale semantic patterns.

High-PepBinder contains 17 Transformer blocks, consisting of 8 encoder layers, 1 bottleneck layer, and 8 decoder layers. Each block follows the standard DiT formulation:

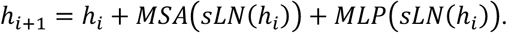

The noisy input representation 𝑥_𝑡_ is transformed into the initial hidden state ℎ_0_ through linear projection and positional encoding, and is then combined with timestep embeddings and protein-conditioning embeddings. During the encoder stage, the hidden state is updated sequentially through the Transformer blocks 𝑖 = 1, …, 8.

During the decoder stage, intermediate encoder features 𝑠_𝑖_ are concatenated with the current decoder state to form skip connections that mitigate information degradation in deep diffusion processes. The update rule for each decoder layer is given by:

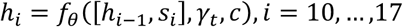

These semantic skip connections enable the model to retain local features captured in early layers (such as local amino-acid motifs and sequence scaffolds) while allowing deeper attention layers to aggregate global contextual information, including long-range dependencies and protein-peptide interaction semantics. This improves conditional-consistent reverse-diffusion denoising. Finally, the decoder produces the noise estimate 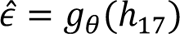 through an output projection.

##### AdaLN-Based Conditional Modulation

To ensure temporal consistency in the diffusion process and effective protein-peptide conditional control, High-PepBinder adopts an adaptive normalization mechanism (AdaLN) within each DiT block, where dynamic scaling and shifting parameters generated from the conditioning vectors are used to modulate Transformer activations. Specifically, let ℎ denote the hidden representation at the current layer, 𝛾_𝑡_ the diffusion timestep embedding, and 𝑐 the protein conditioning embedding. The combined control signal is defined as 𝑧 = 𝛾_𝑡_ + 𝑐. To jointly modulate both the attention and feed-forward pathways, AdaLN applies a multi-head nonlinear mapping to 𝑧, producing six sets of modulation parameters:

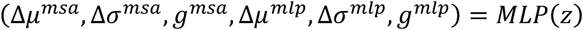

where Δ𝜇 and Δ𝜎 denote the offsets of the mean and variance, respectively, and 𝑔 represents the channel-wise gating coefficient.

The outputs of the multi-head self-attention and MLP sublayers are updated sequentially as follows:

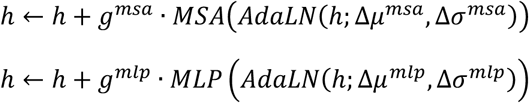

This two-stage modulation with dual gating allows the model to allocate distinct generation strategies according to the diffusion stage and protein conditioning. Early diffusion stages emphasize global structural consistency, whereas later stages focus on fine-grained pattern reconstruction, such as side-chain preferences at key positions and motif-level constraints. Notably, the AdaLN weights are initialized to zero, such that the model initially behaves as a standard LayerNorm-based Transformer and gradually learns conditional modulation. This progressive transition from unconditional to conditional generation has been shown to improve training stability and efficiency by avoiding premature reliance on conditioning signals, and is particularly well suited for conditional generation tasks in biological sequences.

#### 2.1.3 Decoder

Because ESM-C is a pure encoder architecture and does not provide a decoding head, we implement a linear projection decoder consistent with that of ESM-3 to map the latent representations reconstructed by the diffusion model back into the amino-acid token space, enabling end-to-end sequence generation^26^:

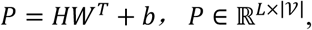

where 𝐻 denotes the diffusion output and 𝑉 represents the amino-acid vocabulary.

#### 2.1.4 Affinity Classifier

We incorporate a lightweight affinity prediction module into High-PepBinder that combines a dual-encoder architecture with a cross-attention mechanism (Figure 1d). The module separately encodes peptides and proteins: the protein branch uses the original ESM-C weights, while the peptide branch uses the ESM-C encoder that has already been finetuned within the diffusion framework. The cross-attention mechanism then explicitly models the dependencies between the peptide and protein representations. For peptide embeddings 𝐻_pep_ and protein embeddings 𝐻_prot_, the cross-attention operation is defined as:

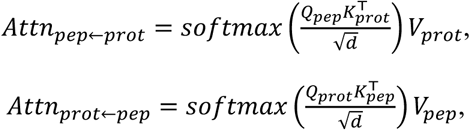

where 𝑄, 𝐾, and 𝑉 denote the query, key, and value matrices obtained through linear projections. After several layers of interaction, the model performs pooling on the sequence-level representations of both the peptide and the protein, concatenates the pooled vectors, and feeds them into an MLP to perform binary affinity classification.

#### 2.1.5 Joint optimization fine-tuning

After the generator and affinity classifier are individually trained, we introduce a lightweight joint optimization fine-tuning phase to gently feedback the affinity discriminative signal into the generative model. Specifically, we update both the diffusion generator 𝐺_𝜃_ and the affinity classifier 𝐶_𝜙_ with a small learning rate, ensuring the generation distribution remains stable while weakly guiding the generation process toward the affinity target.

Additionally, to improve the classifier’s stability under diffusion noise perturbations, we introduce a noise-consistent auxiliary loss: during different diffusion steps t, the generated latent variables are fed as additional inputs to the classifier, constraining the classifier’s outputs to remain consistent under these disturbances. This regularization term helps enhance the classifier’s robustness and indirectly strengthens the consistency of the generation-screening process.

### 2.2 Dataset

#### 2.2.1 Pre-train Dataset

We performed pretraining using a large-scale pseudo protein-peptide dataset derived from Rosetta-based interface fragment extraction, which has been previously employed for training PepInter^42^. The original dataset was constructed by extracting representative linear interface fragments from experimentally resolved protein-protein interaction interfaces to approximate protein-peptide interaction scenarios. It comprises approximately 46 million pseudo protein-peptide pairs, each annotated with an interface energy score. Because Rosetta Peptiderive extracts interface fragments using a sliding-window strategy with a step size of one, the resulting fragment sequences exhibit substantial sequence similarity. To reduce redundancy and mitigate training bias introduced by highly similar samples, we further applied a deduplication procedure to weaken sequence correlations among fragments. In addition, we imposed a threshold on the Rosetta-computed interface energy scores, retaining only samples with interface energies better than -10 Rosetta Energy Units (REU). After these processing steps, we obtained a final set of 8,983,271 high-quality Rosetta-generated pseudo protein-peptide pairs for use in the pretraining stage.

#### 2.2.2 PepPBA Dataset

Due to the scarcity of protein-peptide affinity data, we systematically collected literature published between 1972 and 2025 that reported experimentally measured protein-peptide affinity data, and extracted entries containing quantitative affinity measurements to construct the PepPBA dataset. We searched PubMed, Google Scholar, and Scopus using keywords such as “protein-peptide interaction”, “affinity”, “surface plasmon resonance (SPR)”, “bio-layer interferometry (BLI)”, “fluorescence polarization (FP)”, “enzyme activity assay”, and “microscale thermophoresis (MST)”. We further integrated affinity data from PepLand to increase dataset size and target diversity^29^. In total, PepPBA contains approximately 7,000 records. Each entry includes target information (such as UniProt ID, sequence, and availability of PDB structures), peptide sequence information, and corresponding experimental affinity values (including IC₅₀, K_d_, K_i_, and others).

To improve data quality for model training, we applied strict filtering criteria to PepPBA: (1) target sequence length ≤ 1000 amino acids; (2) peptide sequence length restricted to 6-50 amino acids; (3) removal of entries containing noncanonical amino acids (e.g., X, B) in either target or peptide sequences. After filtering, we retained approximately 5,000 high-quality samples, which were used both as part of the fine-tuning dataset and for training the affinity classification module. For label processing, all affinity values (in molar units) were converted into a negative logarithmic scale to ensure unit consistency. Samples with pAffinity ≥ 6 were classified as high affinity (corresponding to micromolar or tighter binding), whereas those with pAffinity < 6 were classified as low affinity. To our knowledge, PepPBA represents one of the largest and most target-diverse curated protein-peptide affinity datasets currently available, providing a robust and high-quality foundation for training High-PepBinder.

The PepPBA dataset serves both as a component of the fine-tuning dataset for the generative model and as the training dataset for the affinity classification model.

#### 2.2.3 Fine-tuning Dataset

The fine-tuning dataset was constructed by combining the PepPBA dataset described in Section 2.2.2 with protein-peptide interaction data curated from published PDB entries. After removing redundant records, the final fine-tuning dataset comprised 16,380 samples, which were used to fine-tune the generative model the joint optimization stage.

## 3. Results

### 3.1 Evaluating the Target-Specific Affinity Classifier in High-PepBinder

In the protein-peptide affinity prediction task, we systematically evaluated the affinity classification module within High-PepBinder. On the validation set, the classifier achieved an accuracy (ACC) of 0.824, together with strong performance on threshold-independent metrics, including an Area Under the ROC Curve (AUC) of 0.869 and a PR-AUC of 0.807, indicating robust discrimination capability under class imbalance. The F1-score reached 0.681, reflecting a balanced trade-off between precision and recall. Collectively, these results demonstrate that the classifier can reliably distinguish between high- and low-affinity samples. To assess the importance of explicit sequence-interaction modeling, we constructed an ablated model in which the peptide-protein cross-attention mechanism was removed and the two sequences were simply concatenated. Its overall performance decreased by approximately 2 percentage points, confirming that explicit interaction modeling is critical for accurate affinity discrimination.

We also applied Optuna to perform a systematic hyperparameter search for the traditional machine-learning method XGBoost, ensuring that it reached optimal performance under the same input representations^43,44^. Although a tuned XGBoost model achieved comparable results to the deep classifier on some isolated metrics, its performance fluctuated substantially across different targets and distribution settings. This instability arises because traditional ML methods rely on fixed input features, lack the ability to learn hierarchical semantics and conditional dependencies between sequences, and cannot share latent space representations with the generative model. As a result, their generalization to distribution-shifted scenarios is significantly weaker than that of deep representation-based approaches. Considering performance, stability, and framework extensibility, we ultimately selected the deep-learning classifier as the affinity prediction component of High-PepBinder.

We further analyzed latent-space consistency under different training strategies. When trained independently, the classifier achieved an ACC of 0.838. After jointly finetuning it with the generator using a small learning rate and incorporating the noise-consistent auxiliary loss, the final ACC slightly decreased to 0.824. Despite this minor reduction in individual metrics, joint training substantially improved the alignment between the generative and discriminative latent spaces, enhancing robustness on out-of-distribution samples and producing a more coherent, better-generalizing aligned generative-discriminative manifold.

In summary, the affinity prediction module of High-PepBinder strikes a strong balance among accuracy, generalization, and system-level synergy. It provides essential support for high-quality peptide candidate screening and strengthens the end-to-end optimization capabilities of the entire sequence modeling framework.

### 3.2 Sequence and Structural Quality Assessment of Generated Peptides

#### 3.2.1 Evaluation Setup

An effective generative model should be capable of producing diverse candidate structures while remaining consistent with the underlying target distribution. To comprehensively evaluate High-PepBinder’s performance in peptide binder generation, we first conducted systematic tests on the top 30 targets (ranked by sample count) in the PepPBA dataset. This dataset was preprocessed using MMseqs2 for redundancy removal to ensure broad applicability across different targets. For each target protein, we generated 100 candidate peptide sequences for downstream analysis.

The generated peptides were evaluated along two complementary dimensions. First, we assessed the intrinsic quality of the peptide sequences, including sequence diversity, sequence novelty, structural diversity, structural novelty, structural plausibility (the peptide pLDDT calculated by AlphaFold3), amino-acid composition, and latent-space distribution. These metrics allow us to determine whether the model can produce realistic, diverse, and biophysically meaningful peptide sequences. Second, we evaluated the binding capability of the generated peptides to their respective targets using multiple complex-level structural and energetic metrics, including AlphaFold3’s ipTM, ipAE and Rosetta’s 𝑑𝐺_𝑠𝑒𝑝𝑎𝑟𝑎𝑡𝑒𝑑/𝑑𝑆𝐴𝑆𝐴 × 100, to characterize the model’s overall ability to jointly model structure and function in binder design. Definitions of all the above metrics are provided in Section S2 of the SI.

For baseline comparisons, we benchmarked High-PepBinder against state-of-the-art peptide design models, including RFdiffusion and PepMLM^20,21^. RFdiffusion, a leading structure-conditioned generative model, provides a basis for contrasting High-PepBinder with structure-driven frameworks; PepMLM, an advanced sequence language model, enables comparison with purely sequence-based generative paradigms. By evaluating High-PepBinder against both structural and sequence-based models, we provide a more comprehensive characterization of its strengths and advantages in peptide generation.

#### 3.2.2 Sequence- and Structure-Level Generative Properties

We compared two different models in terms of both sequence-level and structure-level diversity and novelty. As shown in Figure 2a, in sequence-level diversity analysis, High-PepBinder consistently achieved the highest sequence divergence across thresholds from 0.2 to 0.5. Notably, at a threshold of 0.4, High-PepBinder maintained a diversity score close to 0.9, whereas RFdiffusion and PepMLM reached only approximately 0.070 and 0.046, respectively, indicating that High-PepBinder can explore a substantially richer sequence space. For sequence novelty, RFdiffusion held a slight advantage, but High-PepBinder still markedly outperformed PepMLM, suggesting that High-PepBinder preserves reasonable exploratory capacity without suffering from mode collapse caused by overly strong structural priors (Figure 2b).

At the structural level, High-PepBinder and PepMLM achieved comparable structural diversity, both significantly higher than RFdiffusion, suggesting that these models are better able to cover a wide range of conformational variations (Figure 2c). In Figure 2d, we observed that structural novelty scores were similar across the three models, indicating that all are capable of generating conformations that differ meaningfully from the training structures. Overall, High-PepBinder demonstrated a balanced and robust generative capability across the four metrics of sequence and structure diversity/novelty, with particularly strong advantages in diversity-related indicators.

Figure 2g compares the Global Amino-acid Composition Discrepancy (GACD) of these three models and the test set. Among them, High-PepBinder produced a distribution most similar to the test set, with a Kullback-Leibler (KL) divergence of 0.0491. PepMLM followed (KL=0.0498), whereas RFdiffusion (KL=0.3504) exhibited a strong bias toward generating alanine (ALA), glutamate (GLU), and leucine (LEU). It is worth noting that PepMLM frequently produced numerous “X” tokens during generation. We filtered out these invalid sequences. without such filtering, the presence of invalid positions would further increase its divergence from the true distribution, meaning the actual KL divergence of PepMLM would be even higher.

For structural foldability assessment, we predicted peptide-protein complexes using AlphaFold3 and extracted peptide pLDDT values as a measure of folding feasibility. As shown in Figure 2f, peptides generated by High-PepBinder achieved an average pLDDT of 57.75, higher than PepMLM (56.63) but slightly lower than RFdiffusion (62.32). Consistent with secondary-structure analysis (Figure 2e), High-PepBinder-generated peptides predominantly adopted loop conformations similar to those of reference peptides, whereas RFdiffusion strongly favored helical structures-an observation consistent with prior studies. Thus, although High-PepBinder generates fewer helices than RFdiffusion, it maintains good structural foldability while adhering to natural conformational statistics, indicating that its generated peptides are structurally plausible.

Finally, latent-space analysis based on ESM embeddings and UMAP revealed clear differences in sequence-manifold organization across models^45^ (Figure 2h-2j). Generated peptides showed substantial overlap with reference peptides and exhibited a more compact and structured manifold, suggesting that High-PepBinder more effectively learns and preserves the evolutionary statistical patterns of natural peptides. In contrast, RFdiffusion and PepMLM displayed more dispersed manifolds with a greater number of outliers deviating from natural sequence distributions.

**Figure 2.**
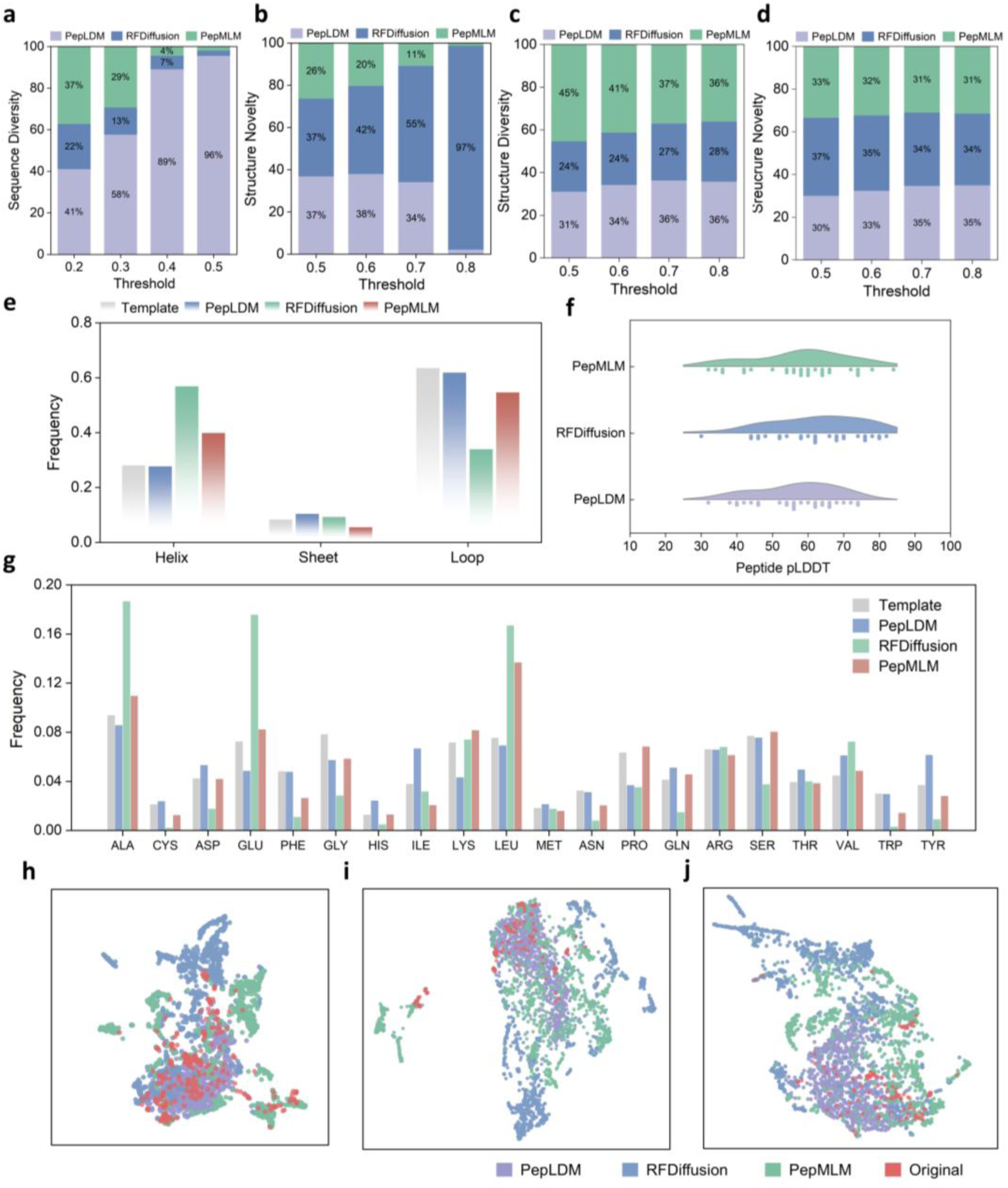
Comprehensive evaluation of High-PepBinder’s generative quality for peptide design. (a-d) Comparison of sequence- and structure-level generative quality among High-PepBinder, RFdiffusion, and PepMLM, including sequence diversity, sequence novelty, structural diversity, and structural novelty. (e) Secondary structure distributions of reference peptides and generated peptides. (f) Average peptide pLDDT scores of generated peptides. (g) GACD between reference peptides and generated peptides. (h-j) Latent-space visualizations based on ESM peptide sequence embeddings using UMAP. To avoid clutter from overlaying all 30 target-specific peptide sets in a single plot, results are shown separately by target groups: (h) first 10 targets; (i) middle 10 targets; (j) final 10 targets, enabling clearer visualization of distribution patterns and model differences.

#### 3.2.3 Binding-Relevant Evaluation: Structural and Energetic Assessment of Generated Peptides

To evaluate the binding capability of the generated peptides to their target proteins, we first used AlphaFold3’s ipTM_mean and ipTM_max scores as structure-level indicators of global complex confidence. High-PepBinder achieved the strongest performance among the three models, with an ipTM_mean of 0.558 across the 30 targets (Figure 3d) and an ipTM_max of 0.851 (Figure 3c). We further defined a “successful hit” as the proportion of generated peptides whose ipTM score exceeds that of the reference peptide for the same target, providing a direct measure of binding advantage at the interface. Figure 3a shows High-PepBinder achieved an average hit rate of 0.200 across all 30 targets, outperforming RFdiffusion (0.175) and PepMLM (0.159). On several targets, this advantage was particularly pronounced. For example, High-PepBinder reached a hit rate of 82% on target 13, demonstrating strong cross-target robustness. We also computed ipAE as an indicator of interface-level conformational reliability, using the median to reduce sensitivity to extreme values (Figure 3e). High-PepBinder again achieved the best performance in ipAE_median across the 30 targets. To further validate binding capability from an energetic perspective, we used Rosetta’s 𝑑𝐺_𝑠𝑒𝑝𝑎𝑟𝑎𝑡𝑒𝑑/𝑑𝑆𝐴𝑆𝐴 × 100 as a complementary metric and quantified the fraction of high-affinity conformations with scores below -1.5 Å². High-PepBinder achieved an average proportion of 0.298, slightly higher than RFdiffusion (0.296) and PepMLM (0.275), indicating a modest yet consistent energetic advantage (Figure 3b).

All in all, High-PepBinder demonstrates superior performance across structural prediction metrics and energy-based evaluations. These results show that, even without explicit structural information and relying only on sequence-level conditioning, High-PepBinder can robustly generate high-quality, target-specific peptide binders, highlighting its strong reliability and practical potential compared with existing methods.

**Figure 3.**
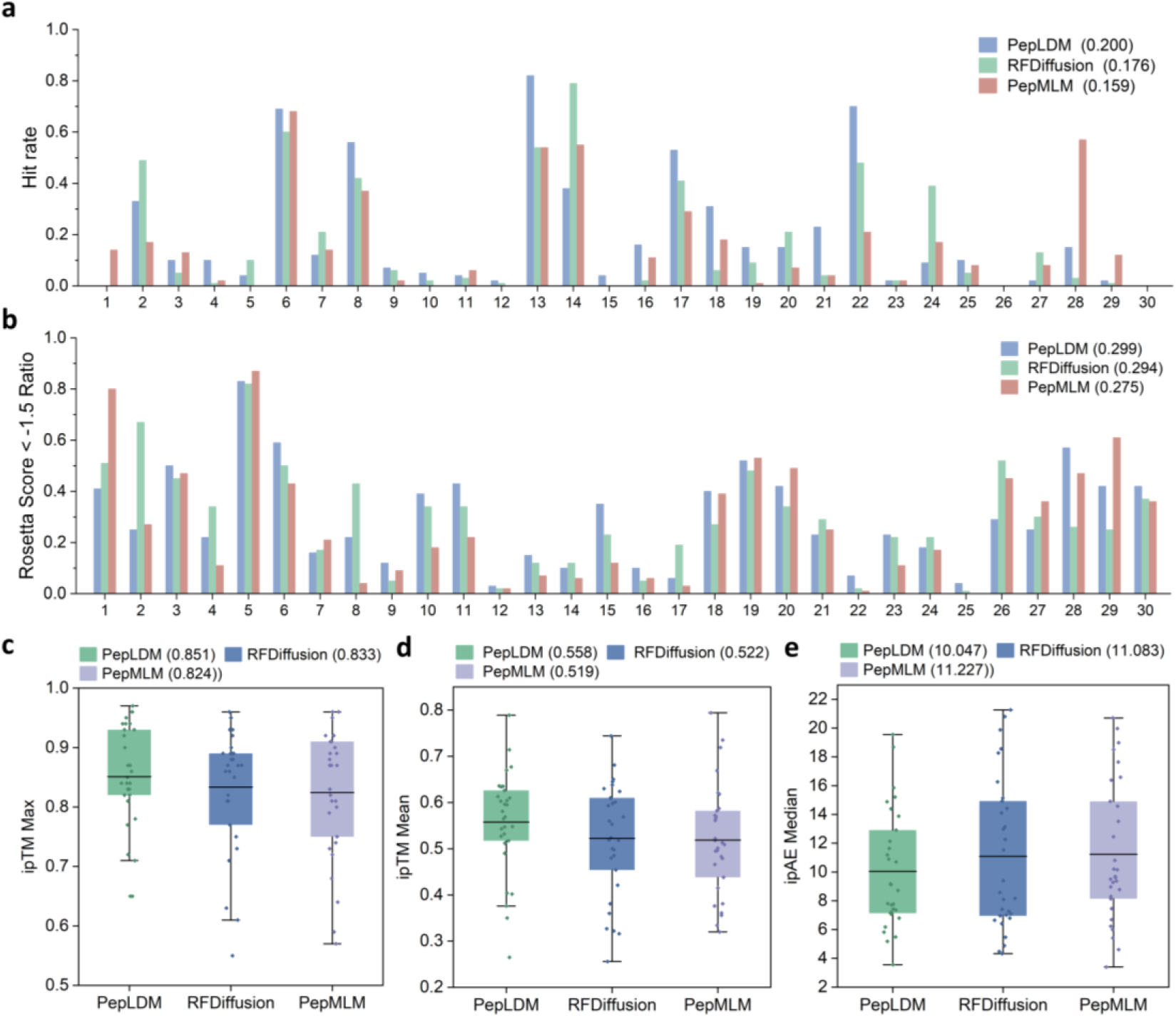
Performance evaluation of High-PepBinder in modeling peptide-protein binding capability. (a) Hit rate of the three models, defined as the proportion of generated peptides whose ipTM exceeds that of the reference peptide. (b) Proportion of generated peptides with Rosetta dG_separated/dSASA×100 < -1.5 Å² for each model. (c) Maximum ipTM achieved by generated peptides for each target across the three models. (d) Mean ipTM of generated peptides for each target. (e) Median ipAE of generated peptides for each target.

### 3.3 De novo peptide binder design using High-PepBinder

We further applied High-PepBinder to generate de novo peptide binders for several important therapeutic targets, namely KEAP1, XIAP, and EGFR. These targets span key biological processes, including oxidative stress response and tumor metabolism regulation (KEAP1), apoptosis control (XIAP) and kinase-driven oncogenic signaling (EGFR). At the same time, they exhibit distinct characteristics in peptide-mediated regulation, ranging from peptide-friendly targets to challenging targets lacking established peptide drug precedents. In addition, to further evaluate the applicability of High-PepBinder across diverse mechanisms of action and target classes, we included several representative targets in supplementary case studies. The corresponding results are provided in the SI. Together, these targets provide a hierarchical and comprehensive testbed for systematically evaluating High-PepBinder’s modeling capability.

For each target, we used AlphaFold3 to predict the structures of candidate peptides with the target protein, followed by unified filtering based on structural confidence metrics and Rosetta interface energies. We then retained only those peptides that occupied the same binding pocket as the reference peptide and selected top 3 candidates with the highest overall scores for subsequent analysis (more details can be found in Section S3 of the SI).

**Figure 4.**
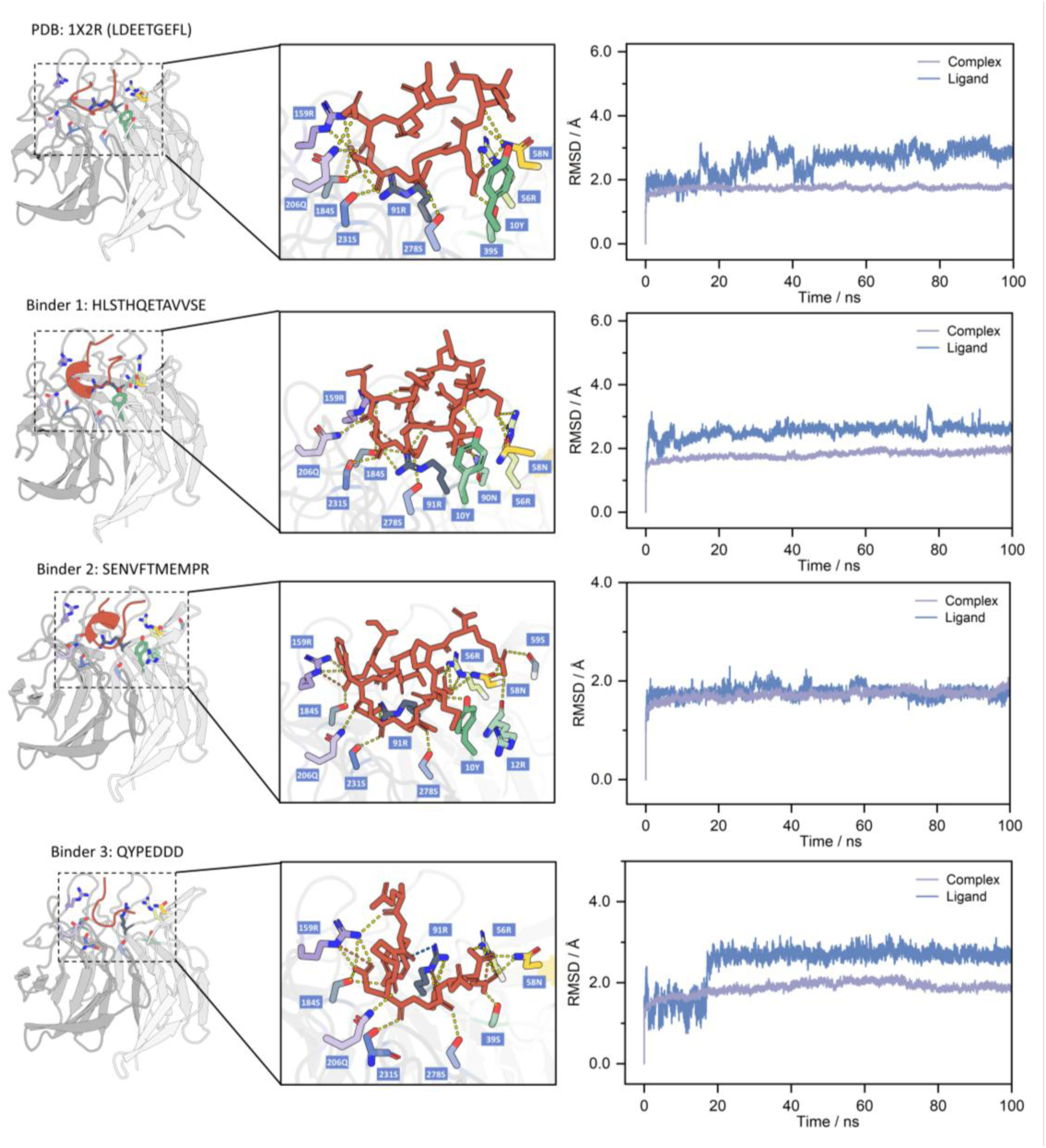
Peptide generation results for the KEAP1 target. Structure 1X2R represents the reference complex between KEAP1 and Nrf2. Binders 1-3 denote candidate peptides generated by High-PepBinder. The left panel shows the overall complex structures and schematic illustrations of secondary interactions at the binding site, where yellow indicates hydrogen bonds, orange indicates salt bridges, and blue indicates cation-π interactions. The right panel presents the root-mean-square deviation (RMSD) of the complexes and ligands during molecular dynamics simulations, which is used to assess binding stability.

**Table 1.**
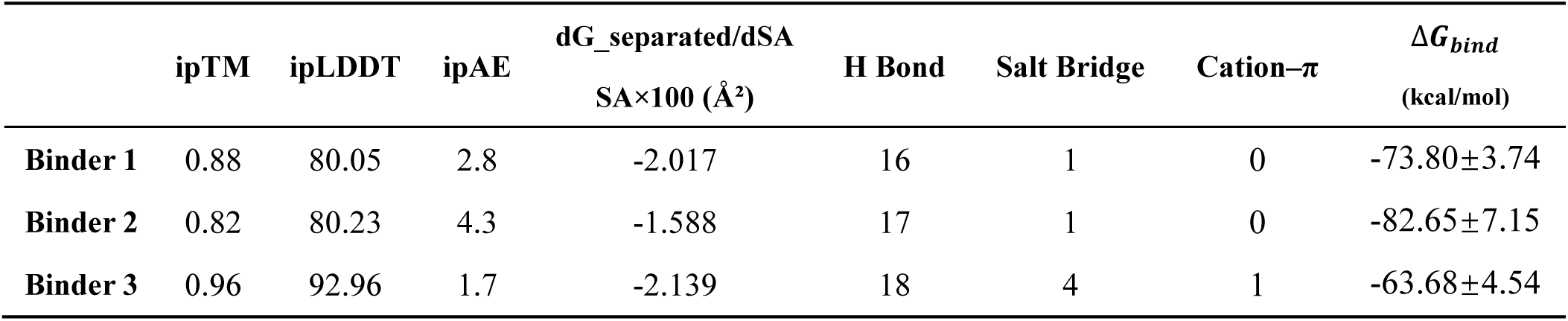
The reference complex 1X2R exhibits a binding free energy (Δ𝐺_𝑏𝑖𝑛𝑑_) of -69.79±4.22 kcal/mol, forms 15 hydrogen bonds, and shows no detectable salt bridges or cation-π interactions.

| | ipTM | ipLDDT | ipAE | dG_separated/dSA<br>SA$\times 100$ ($\text{\AA}^2$) | H Bond | Salt Bridge | Cation-π | $\Delta G_{bind}$<br>(kcal/mol) |
| --- | --- | --- | --- | --- | --- | --- | --- | --- |
| **Binder 1** | 0.88 | 80.05 | 2.8 | -2.017 | 16 | 1 | 0 | $-73.80 \pm 3.74$ |
| **Binder 2** | 0.82 | 80.23 | 4.3 | -1.588 | 17 | 1 | 0 | $-82.65 \pm 7.15$ |
| **Binder 3** | 0.96 | 92.96 | 1.7 | -2.139 | 18 | 4 | 1 | $-63.68 \pm 4.54$ |

Kelch-like ECH-associated protein 1 (KEAP1) is a key regulator of the cellular oxidative stress response, mediating the ubiquitination and degradation of the transcription factor NRF2 through recognition of short linear peptide motifs. The KEAP1-NRF2 axis plays critical roles in tumor initiation and progression, therapy resistance, and metabolic reprogramming, and has therefore long been a focus of drug discovery efforts. Structural analysis of the predicted complexes shows that the KEAP1 residues contacted by the generated peptides overlap by approximately 90% with the reference peptide (PDB: 1X2R), suggesting that the major geometric features of KEAP1 recognition of linear peptide motifs are preserved in the predicted structures (Figure 4)^46^. Building on this conserved binding geometry, the candidate peptides introduce additional interfacial interactions. In particular, candidate binder 3 (QYPEDDD) forms four additional salt bridges and one cation-π interaction between the Tyr2 of the peptide and Arg91 of KEAP1. In contrast, only a hydrogen-bond interaction is observed at the corresponding position in the reference complex. Molecular dynamics simulations and binding free energy analyses further show that all three candidate peptides exhibit binding free energies comparable to or lower than that of the reference complex under identical geometric constraints. (Table 1; see Section S4 of the SI for simulation details). Together, these results indicate favorable predicted interfacial stability of the High-PepBinder-generated peptides relative to the reference system, supporting their potential as candidate binders.

**Figure 5.**
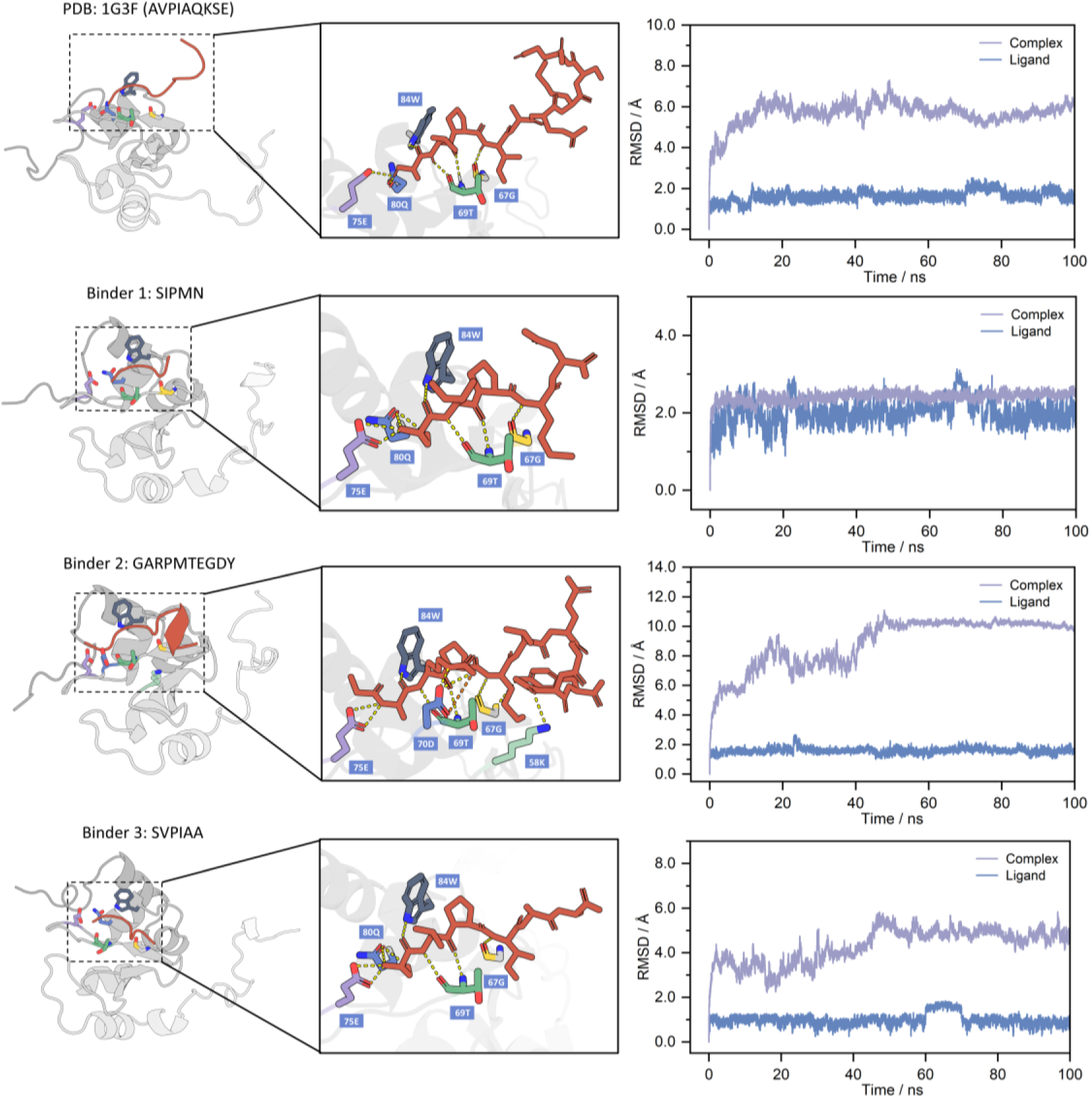
Peptide generation results for the XIAP target. Structure 1G3F represents the reference complex between the XIAP BIR3 domain and the SMAC nonapeptide. Binders 1-3 denote candidate peptides generated by High-PepBinder.

**Table 2.** The reference complex 1G3F exhibits a binding free energy (Δ𝐺_𝑏𝑖𝑛𝑑_) of -45.88±4.44 kcal/mol, forms 6 hydrogen bonds, and shows no detectable salt bridges or cation-π interactions.

| | ipTM | ipLDDT | ipAE | dG_separated/dSA<br>SA$\times 100$ ($\text{\AA}^2$) | H Bond | Salt Bridge | $\Delta G_{bind}$<br>(kcal/mol) |
| --- | --- | --- | --- | --- | --- | --- | --- |
| **Binder 1** | 0.84 | 82.80 | 2.7 | -3.167 | 8 | 0 | $-49.53 \pm 4.40$ |
| **Binder 2** | 0.83 | 82.18 | 2.9 | -1.990 | 10 | 1 | $-62.45 \pm 3.25$ |
| **Binder 3** | 0.83 | 80.17 | 2.6 | -3.275 | 8 | 0 | $-50.52 \pm 4.67$ |

The X-linked inhibitor of apoptosis protein (XIAP) is a key negative regulator of the apoptotic pathway, where it blocks programmed cell death by inhibiting caspase activation. Owing to its aberrant overexpression in multiple cancers, XIAP is widely regarded as a therapeutically relevant antitumor target. The representative candidate binder 3 (SVPIAA) adopts a spatial conformation similar to that of the reference SMAC nonapeptide (PDB: 1G3F) in the predicted complex, and its sequence features are consistent with established preferences of the BIR3 pocket, where an N-terminal hydrophobic residue in combination with proline promotes a compact conformation and precise hydrophobic anchoring^47^ (Figure 5). Molecular dynamics simulations further indicate that the binding free energies of the candidate peptide-XIAP complexes are lower than that of the reference system under identical simulation protocols. Despite differences in sequence composition, the candidates display consistent pocket occupancy and key residue interaction patterns at the structural level, suggesting that High-PepBinder can generate sequence-diverse peptide candidates with favorable predicted binding characteristics.

**Figure 6.**
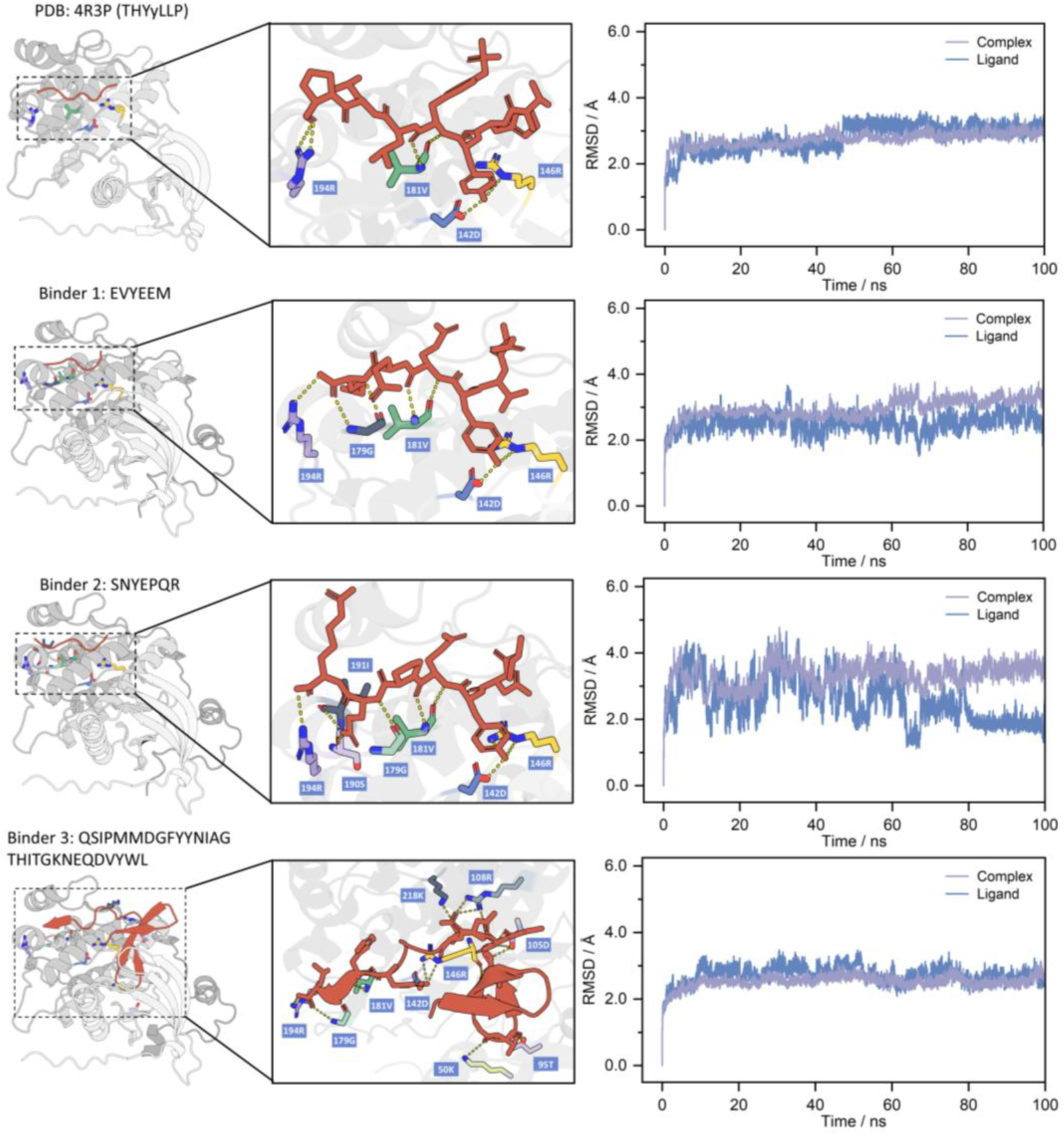
Peptide generation results for the EGFR target. Structure 4R3P represents the reference complex between the EGFR domain and the Mig6 peptide. Binders 1-3 denote candidate peptides generated by High-PepBinder.

**Table 3.** The reference complex 4R3P exhibits a binding free energy (𝛥𝐺*_bind_*) of -53.67±3.03 kcal/mol, forms 6 hydrogen bonds, and shows no detectable salt bridges or cation-π interactions.

| | ipTM | ipLDDT | ipAE | dG_separated/dSA<br>SA$\times 100$ ($\text{\AA}^2$) | H Bond | $\Delta G_{bind}$<br>(kcal/mol) |
| --- | --- | --- | --- | --- | --- | --- |
| **Binder 1** | 0.92 | 82.90 | 3.0 | -2.119 | 8 | $-50.80 \pm 3.70$ |
| **Binder 2** | 0.92 | 82.98 | 2.8 | -2.506 | 9 | $-58.76 \pm 3.67$ |
| **Binder 3** | 0.87 | 80.26 | 4.0 | -2.046 | 17 | $-149.40 \pm 6.63$ |

The epidermal growth factor receptor (EGFR) is a well-studied oncogenic kinase and a challenging target for peptide-based modulation. We used the MIG6-bound EGFR complex (PDB: 4R3P) as a reference to evaluate peptide generation at a non-ATP regulatory interface^48^. As summarized in Table 3, the generated peptide candidates exhibit high predicted complex confidence, low interface prediction error, and favorable calculated interfacial energetics, including normalized interface energies and binding free energies comparable to or lower than the reference system. Together, these results suggest that High-PepBinder can generate peptide candidates with consistent predicted structural and energetic properties at the EGFR regulatory interface under identical simulation protocols.

## 4. Conclusion

With the rapid advancement of generative models and pLMs in biological sequence modeling, de novo design of target-specific peptides has become feasible^49,50^. However, existing peptide generation methods either rely on explicit 3D structural information or lack direct constraints on functional objectives, making it difficult to achieve stable and controllable design for targets with missing or highly dynamic structures^19–21,49,51^. This tension highlights the urgent need for a unified peptide design framework that can simultaneously enable target-conditioned control and functional optimization under structure-independent settings.

In this study, we propose High-PepBinder, a peptide design framework that integrates pLMs, latent diffusion-based generation, and affinity-aware modeling within a unified latent space. Through systematic benchmark evaluations and representative target-centered case studies, High-PepBinder demonstrates robust performance in generating target-conditioned peptide candidates with favorable predicted structural and energetic properties. In AlphaFold 3-based structural assessments, High-PepBinder achieves an average “successful hit rate” of approximately 20% across 30 targets, with substantially higher hit rates observed for certain targets, indicating strong generalization across diverse target conditions. For multiple therapeutic targets, including KEAP1, XIAP, and EGFR, the generated peptides recapitulate key interfacial features observed in the reference systems within the predicted complexes, while exhibiting increased sequence diversity and comparable or more favorable predicted binding free energies under identical computational settings.

From a methodological perspective, High-PepBinder establishes a structure-independent peptide design paradigm by jointly modeling conditional generation and affinity-aware screening within a unified framework and incorporating affinity information through two complementary mechanisms. This design alleviates key limitations of traditional peptide engineering approaches and provides a general computational strategy for peptide discovery that complements structure-driven methods, particularly for challenging targets.

Although High-PepBinder has demonstrated encouraging performance, several directions remain for further exploration. Future work could expand the affinity dataset to cover a broader range of target classes, thereby improving model robustness in data-scarce settings. In addition, peptide therapeutics often require optimization beyond binding affinity, including properties such as membrane permeability and plasma stability. Incorporating multi-objective optimization or reinforcement learning into High-PepBinder’s generative process may further enhance the drug development potential of the generated peptides^52^.

Overall, High-PepBinder provides a general, flexible, and forward-looking computational framework for peptide drug discovery. We believe that, with continued improvements in data quality and model capability, this framework has the potential to play an important role in future targeted therapeutics and functional peptide engineering.

## Data Availability Statement

The data that support the findings of this study are available from the corresponding author upon reasonable request. The data that support the findings of this study are available from the corresponding author upon reasonable request.

## Supporting Information

The complete raw data include: (1) supplementary descriptions of the datasets; (2) detailed definitions and full results of the High-PepBinder performance evaluation module, covering sequence novelty, sequence diversity, structural novelty, structural diversity, structural plausibility, ipTM, ipAE, and Rosetta’s dG_separated/dSASA×100; (3) the screening pipeline for the case studies, along with molecular dynamics model construction and binding free energy calculation workflows; and (4) additional case-study targets that differ from those in the main text in both mechanism of action and protein class, including the nuclear receptor ESRRA and the E3 ubiquitin ligase adaptor TRAF6, which are used to further assess High-PepBinder’s generalization across diverse target types.

## Acknowledgements

This project was supported by Macao Science and Technology Development Fund (Grant No. 0029/2025/RIB1) and the internal grant from Macao Polytechnic University (RP/FCA-11/2025).

## Author contributions

These authors contributed equally to this work: Q. M., S. Z. Conceptualization and investigation: Q. M., S. Z., H. D. Data acquisition: Q. M., W. X., N. Z. Develop the model: Q.M., S.Z. Visualized data and created figures: Q. M. Writing-original draft: Q. M. Writing-review & editing: Q. M. Funding acquisition: H. D. Project administration: J. G., H. D.

