## Supporting information for "High-PepBinder: A pLM-Guided Latent Diffusion Framework for Affinity-Aware Target-Specific Peptide Design"

### Section S1. PepInter Dataset

The pseudo protein-peptide dataset and the protein-peptide interaction dataset were derived from the PepInter study, and the construction procedures for the two datasets are as follows[1]:

#### *1. Pseudo protein-peptide dataset*

The pseudo protein-peptide complex dataset was derived from the PPI3D database, which contains more than 891,000 experimentally determined protein-protein interaction (PPI) pairs with resolved three-dimensional structures. To ensure data reliability and robustness, several filtering criteria were applied during preprocessing: (i) structural resolution better than 4.0 Å; (ii) protein sequence length shorter than 1000 amino acids; and (iii) removal of PPI pairs with sequence identity greater than 40% to reduce redundancy. MMseqs2 was employed to cluster all protein sequences using a 40% sequence identity threshold. After filtering and clustering, a total of 28,916 non-redundant PPI pairs with experimentally resolved PDB structures were retained.

Subsequently, the Rosetta Peptiderive protocol was applied to these PPIs to generate pseudo protein-peptide fragment pairs. For each protein-protein complex, one chain was fixed as the receptor protein, while the interacting partner chain was systematically decomposed into linear peptide fragments using a sliding-window strategy with fragment lengths ranging from 5 to 50 amino acids. For each generated pseudo peptide-protein pair, Peptiderive computed an interface energy score to quantify the energetic contribution of the peptide fragment to the interaction interface. Pairs associated with abnormal or erroneous interface scores were discarded. Finally, approximately 46 million pairs of data samples were generated.

#### *2. Protein-peptide interaction dataset*

The experimentally validated protein-peptide interaction dataset was constructed by systematically scanning all protein complex structures deposited in the RCSB Protein Data Bank (PDB, accessed in September 2025). A Python-based pipeline was developed to identify interacting chain pairs, where two chains were considered to form an interaction if any pair of heavy atoms between them was within a distance threshold of 5 Å. To specifically capture protein-peptide interactions, interacting chain pairs were further filtered by chain length, requiring the protein chain to contain 40-1000 residues and the peptide chain to contain 5-40 residues. Redundant interaction pairs with identical protein-peptide sequence combinations were removed. Following this procedure, a total of 11,380 non-redundant protein-peptide interaction pairs were obtained.

### Section S2. Evaluation Metrics for Peptide Generation Quality

To evaluate the quality of peptides generated by High-PepBinder, we use the following evaluation metrics:

#### *1. Sequence Diversity and Novelty*

We computed sequence diversity and novelty metrics for the generated peptides using the PairwiseAligner module in Biopython with the BLOSUM62 substitution matrix and a global alignment scheme. For any two sequence $x$ and $y$, we define their similarity as:

$$sim(x, y) = \frac{S(x, y)}{\sqrt{S(s, y)S(y, y)}}$$

where $S(\cdot, \cdot)$ denotes the alignment score obtained under the same alignment parameters. The sequence distance is then defined as $1 - sim(x, y)$. Based on the resulting distance matrix, we performed hierarchical clustering and partitioned all generated peptides into clusters using a threshold of $t$. The diversity score is defined as the ratio between the number of clusters and the total number of generated peptides. For novelty, a generated peptide $g$ is considered novel if its maximum similarity to any reference peptide $o$ satisfies $max_o sim(g, o) \leq 1 - t$. The overall novelty score is computed as the proportion of generated peptides classified as novel.

#### 2. *Structure Diversity and Novelty*

We computed structural similarity between generated peptides using the TM-score, and defined the structural distance as $1 - TM^2$. We then applied hierarchical clustering with a TM-score threshold $t$ to group the generated peptides, and treated the resulting clusters as structural clusters. The structural diversity score is defined as the ratio of the number of clusters to the total number of generated peptides, reflecting the breadth and heterogeneity of the generated conformational space.

In addition, to quantify the structural differences between generated peptides and reference peptides, we computed, for each generated peptide, its maximum TM-score against all reference peptides to characterize structural similarity. The structural novelty of a peptide is then measured as $1 - TM_{best}$.

***3. Structural foldability***

Structural plausibility was quantified using the peptide-level pLDDT predicted by AlphaFold3, where peptide pLDDT > 70 generally indicates a reasonably reliable backbone fold. However, most linear peptides are intrinsically disordered or form only short secondary-structure segments under physiological conditions, and therefore typically exhibit lower pLDDT values.

***4. Interface TM score (ipTM)***

The ipTM score is used to evaluate the reliability of predicted inter-chain arrangements and interface interactions in protein complexes, and serves as an important indicator of complex structural quality.

***5. Rosetta Score***

Rosetta's dG_separated/dSASA×100 metric quantifies the normalized energetic contribution of interface binding relative to changes in solvent-accessible surface area. Here, $dG\_separated$ represents the change in interfacial free energy upon complex dissociation, and $dSASA$ denotes the difference in solvent-accessible surface area before and after binding. This ratio measures the binding energy per unit interface area.

***6. ipAE***

Interface Predicted Alignment Error (ipAE) is used to assess the reliability of AlphaFold 3's predictions in the protein-peptide interface region. Interface residues are defined as residue pairs between the protein and peptide whose minimum interatomic distance is less than 8 Å. From the full PAE matrix, we extract the predicted alignment errors corresponding to these interface residue pairs and compute their median value, denoted as ipAE (median). Smaller ipAE values indicate more reliable local conformational predictions at the interface and more trustworthy relative spatial relationships between the peptide and the protein.

### Section S3. Algorithms for Evaluation Metrics in the Case Study

After generating peptide sequences with High-PepBinder, we applied the following filtering pipeline to identify the top-3 candidates. First, we used AlphaFold3 to predict the structures of peptide-protein complexes and retained only those satisfying all of the following criteria: ipTM > 85, peptide interface-pLDDT (iPLDDT) > 80, ipAE < 5 Å, Rosetta's dG_separated/dSASA × 100 < –1.5Å$^2$ [3-5], and located within the same binding pocket as the reference peptide. Peptides that passed all filtering criteria were then subjected to Amber molecular dynamics (MD) simulations to further assess their binding stability, conformational robustness, and the physical plausibility of their interface interactions[6]. The MD workflow and the calculation methods for each evaluation metric are described below.

#### *1. Peptide interface-PLDDT*

For each predicted protein–peptide complex structure, we first used mdtraj to identify interface residues between the peptide chain (chain B) and the protein chain (chain A) based on the closest heavy-atom distance, using an 8 Å cutoff [7]. Using these interface residues as indices, we extracted the atomic pLDDT values stored in the original B-factor field and computed the mean pLDDT for each residue. Finally, peptide interface-PLDDT was defined as the arithmetic average of the per-residue pLDDT values across all peptide interface residues, providing a measure of the model's confidence and structural stability for the peptide in the interface region.

### Section S4. Molecular Dynamics

#### *1. Molecular Dynamics Simulations*

Molecular dynamics simulations were performed using Amber 24 to evaluate the binding affinity between the screened sequences and the target protein[8,9]. Experimentally resolved target-ligand crystal complexes were used as reference systems, while complexes formed by the screened sequences were generated using the AlphaFold3 structure prediction method. All complexes were solvated in a truncated octahedral box of TIP3P water molecules with a minimum solute-boundary distance of 10.0 Å, and counterions were added to neutralize the systems. The ff14SB force field was employed throughout the simulations.

Energy minimization was carried out in two stages. First, 10,000 steps of minimization were performed with a harmonic restraint of 500 kcal $mol^{-1}$ $Å^{-2}$ applied to the solute to relax solvent molecules and ions, followed by an additional 10,000 steps of unrestrained minimization. The systems were then gradually heated from 0 K to 300 K under the NVT ensemble over 100 ps, during which a weak positional restraint of 10 kcal $mol^{-1}$ $Å^{-2}$ was applied to the solute. After removal of all restraints, production MD simulations were conducted under the NPT ensemble at 300 K and 1 atm for 100-120 ns. Trajectories were saved every 10 ps for subsequent structural analysis and MM/GBSA binding free energy calculations. An identical simulation protocol was applied to all systems to ensure consistency and comparability.

#### 2. *Binding Free Energy Analysis*

The binding free energies between the screened sequences and the target protein were evaluated using the MM/GBSA method. For each complex, 50 snapshots were evenly extracted from a 20 ns stable segment of the MD trajectory for energy calculations. The binding free energy was defined as the free energy difference between the complex and the sum of the receptor and ligand.

In MM/GBSA calculations, the binding free energy consists of gas-phase molecular mechanical energy and solvation free energy. The gas-phase energy includes electrostatic and van der Waals interaction terms, while the solvation free energy comprises polar and nonpolar contributions. The polar solvation energy was estimated using the generalized Born (GB) model with interior and exterior dielectric constants set to 4 and 80, respectively[10,11]. The nonpolar solvation energy was calculated based on the LCPO algorithm using solvent-accessible surface area[12]. The resulting binding free energies were used for relative comparison among different complexes.

### Section S5. Results on the Generative Quality of Peptide Sequences

*1. Peptide pLDDT*

| Target | High-PepBinder | RFDiffusion | PepMLM |
|---|---|---|---|
| 1 | 69.558 | 77.099 | 83.089 |
| 2 | 51.891 | 67.506 | 54.342 |
| 3 | 54.267 | 52.198 | 54.501 |
| 4 | 45.573 | 43.669 | 41.683 |
| 5 | 72.001 | 80.370 | 77.032 |
| 6 | 42.013 | 57.441 | 57.879 |
| 7 | 56.363 | 69.220 | 60.589 |
| 8 | 43.891 | 53.147 | 35.974 |
| 9 | 74.871 | 76.373 | 74.973 |
| 10 | 61.557 | 51.683 | 42.451 |
| 11 | 55.387 | 61.061 | 56.085 |
| 12 | 63.909 | 67.377 | 55.884 |
| 13 | 37.919 | 30.198 | 34.313 |
| 14 | 66.177 | 80.218 | 73.747 |
| 15 | 55.177 | 62.426 | 49.063 |
| 16 | 40.509 | 45.035 | 43.735 |
| 17 | 32.032 | 46.975 | 32.610 |
| 18 | 61.035 | 62.961 | 62.000 |
| 19 | 73.332 | 62.400 | 57.925 |
| 20 | 66.897 | 74.699 | 66.882 |
| 21 | 40.155 | 47.475 | 36.084 |
| 22 | 63.754 | 67.143 | 60.893 |
| 23 | 55.353 | 58.420 | 66.826 |
| 24 | 68.012 | 82.053 | 71.249 |
| 25 | 66.676 | 75.821 | 63.771 |
| 26 | 61.620 | 75.293 | 74.422 |

| | | | |
|---|---|---|---|
| 27 | 54.063 | 66.842 | 64.108 |
| 28 | 46.494 | 44.713 | 58.066 |
| 29 | 58.262 | 59.887 | 59.199 |
| 30 | 60.126 | 69.981 | 63.159 |

**Table S1.** Peptide pLDDT results

#### *2. The Global Amino-acid Composition Discrepancy (GACD)*

| Amino Acid | Template | High-PepBinder | RFDiffusion | PepMLM |
|---|---|---|---|---|
| ALA | 0.094 | 0.086 | 0.186 | 0.109 |
| CYS | 0.021 | 0.024 | 0.002 | 0.012 |
| ASP | 0.042 | 0.053 | 0.017 | 0.042 |
| GLU | 0.072 | 0.048 | 0.175 | 0.082 |
| PHE | 0.048 | 0.047 | 0.011 | 0.026 |
| GLY | 0.078 | 0.057 | 0.028 | 0.058 |
| HIS | 0.013 | 0.024 | 0.005 | 0.013 |
| ILE | 0.038 | 0.067 | 0.032 | 0.020 |
| LYS | 0.071 | 0.043 | 0.074 | 0.081 |
| LEU | 0.075 | 0.069 | 0.167 | 0.137 |
| MET | 0.018 | 0.021 | 0.017 | 0.016 |
| ASN | 0.032 | 0.031 | 0.008 | 0.020 |
| PRO | 0.063 | 0.037 | 0.035 | 0.068 |
| GLN | 0.041 | 0.051 | 0.015 | 0.045 |
| ARG | 0.066 | 0.066 | 0.068 | 0.061 |
| SER | 0.077 | 0.075 | 0.037 | 0.080 |
| THR | 0.039 | 0.049 | 0.040 | 0.038 |
| VAL | 0.045 | 0.061 | 0.072 | 0.048 |
| TRP | 0.030 | 0.029 | 0.003 | 0.014 |
| TYR | 0.037 | 0.061 | 0.009 | 0.028 |

**Table S2.** The Global Amino-acid Composition Discrepancy (GACD) results

#### *3. Distribution of secondary structures*

| Secondary Structure | Template | High-PepBinder | RFDiffusion | PepMLM |
|---|---|---|---|---|
| Helix | 0.2809 | 0.2772 | 0.5686 | 0.3985 |
| Sheet | 0.0836 | 0.1041 | 0.0924 | 0.0552 |
| Loop | 0.6356 | 0.6187 | 0.339 | 0.5464 |

**Table S3.** Distribution of secondary structures

#### *4. Sequence diversity results of High-PepBinder*

| Target | Threshold | | | |
|---|---|---|---|---|
| | 0.2 | 0.3 | 0.4 | 0.5 |
| 1 | 1.00 | 1.00 | 0.98 | 0.52 |
| 2 | 1.00 | 1.00 | 0.96 | 0.62 |
| 3 | 1.00 | 0.98 | 0.90 | 0.43 |
| 4 | 1.00 | 1.00 | 0.97 | 0.54 |
| 5 | 1.00 | 1.00 | 0.99 | 0.69 |
| 6 | 1.00 | 1.00 | 0.93 | 0.54 |
| 7 | 1.00 | 1.00 | 0.97 | 0.67 |
| 8 | 1.00 | 0.98 | 0.93 | 0.52 |
| 9 | 1.00 | 1.00 | 0.96 | 0.53 |
| 10 | 1.00 | 0.98 | 0.82 | 0.22 |
| 11 | 1.00 | 1.00 | 0.87 | 0.24 |
| 12 | 1.00 | 1.00 | 0.99 | 0.65 |
| 13 | 1.00 | 1.00 | 0.98 | 0.59 |
| 14 | 1.00 | 1.00 | 0.97 | 0.69 |
| 15 | 1.00 | 1.00 | 1.00 | 0.71 |
| 16 | 1.00 | 1.00 | 0.95 | 0.61 |
| 17 | 1.00 | 1.00 | 0.98 | 0.64 |
| 18 | 1.00 | 1.00 | 0.93 | 0.55 |

| | | | | |
|---|---|---|---|---|
| 19 | 1.00 | 1.00 | 0.97 | 0.61 |
| 20 | 1.00 | 1.00 | 0.96 | 0.68 |
| 21 | 1.00 | 1.00 | 0.95 | 0.50 |
| 22 | 1.00 | 0.98 | 0.94 | 0.52 |
| 23 | 1.00 | 0.99 | 0.97 | 0.52 |
| 24 | 1.00 | 0.99 | 0.95 | 0.51 |
| 25 | 0.99 | 0.99 | 0.99 | 0.56 |
| 26 | 1.00 | 0.99 | 0.87 | 0.29 |
| 27 | 1.00 | 1.00 | 0.98 | 0.67 |
| 28 | 1.00 | 1.00 | 0.97 | 0.63 |
| 29 | 1.00 | 1.00 | 0.98 | 0.72 |
| 30 | 1.00 | 1.00 | 0.99 | 0.79 |

**Table S4.** Sequence diversity results of High-PepBinder

#### *5. Sequence diversity results of RFDiffusion*

| **Target** | **Threshold** | | | |
|---|---|---|---|---|
| | **0.2** | **0.3** | **0.4** | **0.5** |
| 1 | 0.62 | 0.30 | 0.12 | 0.01 |
| 2 | 0.33 | 0.18 | 0.09 | 0.03 |
| 3 | 0.51 | 0.15 | 0.04 | 0.01 |
| 4 | 0.62 | 0.29 | 0.07 | 0.02 |
| 5 | 0.57 | 0.30 | 0.07 | 0.02 |
| 6 | 0.55 | 0.16 | 0.02 | 0.01 |
| 7 | 0.49 | 0.25 | 0.11 | 0.02 |
| 8 | 0.53 | 0.30 | 0.12 | 0.01 |
| 9 | 0.30 | 0.11 | 0.06 | 0.02 |
| 10 | 0.57 | 0.31 | 0.15 | 0.01 |
| 11 | 0.69 | 0.23 | 0.05 | 0.03 |
| 12 | 0.58 | 0.22 | 0.01 | 0.01 |

| | | | | |
|---|---|---|---|---|
| 13 | 0.72 | 0.32 | 0.04 | 0.01 |
| 14 | 0.66 | 0.29 | 0.07 | 0.02 |
| 15 | 0.39 | 0.13 | 0.02 | 0.01 |
| 16 | 0.67 | 0.26 | 0.08 | 0.01 |
| 17 | 0.43 | 0.11 | 0.04 | 0.01 |
| 18 | 0.57 | 0.18 | 0.01 | 0.01 |
| 19 | 0.42 | 0.19 | 0.06 | 0.01 |
| 20 | 0.52 | 0.25 | 0.07 | 0.01 |
| 21 | 0.61 | 0.35 | 0.13 | 0.02 |
| 22 | 0.61 | 0.22 | 0.02 | 0.01 |
| 23 | 0.38 | 0.21 | 0.10 | 0.01 |
| 24 | 0.37 | 0.16 | 0.07 | 0.01 |
| 25 | 0.53 | 0.22 | 0.14 | 0.02 |
| 26 | 0.53 | 0.22 | 0.15 | 0.03 |
| 27 | 0.61 | 0.22 | 0.01 | 0.01 |
| 28 | 0.27 | 0.12 | 0.02 | 0.01 |
| 29 | 0.69 | 0.29 | 0.07 | 0.01 |
| 30 | 0.47 | 0.25 | 0.08 | 0.02 |

**Table S5.** Sequence diversity results of RFDiffusion

#### *6. Sequence diversity results of PepMLM*

| **Target** | **Threshold** | | | |
|---|---|---|---|---|
| | **0.2** | **0.3** | **0.4** | **0.5** |
| 1 | 0.99 | 0.75 | 0.05 | 0.01 |
| 2 | 0.97 | 0.65 | 0.05 | 0.01 |
| 3 | 0.96 | 0.35 | 0.01 | 0.01 |
| 4 | 0.63 | 0.17 | 0.01 | 0.01 |
| 5 | 1.00 | 0.63 | 0.05 | 0.01 |
| 6 | 0.74 | 0.14 | 0.01 | 0.01 |

| | | | | |
|---|---|---|---|---|
| 7 | 0.91 | 0.23 | 0.01 | 0.01 |
| 8 | 0.79 | 0.38 | 0.02 | 0.01 |
| 9 | 0.99 | 0.46 | 0.02 | 0.01 |
| 10 | 0.96 | 0.41 | 0.01 | 0.01 |
| 11 | 0.88 | 0.37 | 0.01 | 0.01 |
| 12 | 0.91 | 0.75 | 0.04 | 0.01 |
| 13 | 0.93 | 0.51 | 0.02 | 0.01 |
| 14 | 0.41 | 0.02 | 0.01 | 0.01 |
| 15 | 1.00 | 0.79 | 0.08 | 0.01 |
| 16 | 0.89 | 0.38 | 0.01 | 0.01 |
| 17 | 0.97 | 0.77 | 0.14 | 0.01 |
| 18 | 0.93 | 0.67 | 0.09 | 0.01 |
| 19 | 0.96 | 0.43 | 0.02 | 0.01 |
| 20 | 0.99 | 0.69 | 0.09 | 0.01 |
| 21 | 0.93 | 0.66 | 0.17 | 0.02 |
| 22 | 0.98 | 0.86 | 0.11 | 0.01 |
| 23 | 0.94 | 0.78 | 0.08 | 0.02 |
| 24 | 0.95 | 0.39 | 0.02 | 0.01 |
| 25 | 0.77 | 0.04 | 0.01 | 0.01 |
| 26 | 1.00 | 0.90 | 0.07 | 0.01 |
| 27 | 0.96 | 0.58 | 0.04 | 0.01 |
| 28 | 0.93 | 0.43 | 0.03 | 0.01 |
| 29 | 0.92 | 0.61 | 0.05 | 0.01 |
| 30 | 0.87 | 0.35 | 0.04 | 0.01 |

**Table S6.** Sequence diversity results of PepMLM

#### *7. Sequence novelty results of High-PepBinder*

| **Target** | **Threshold** | | | |
|---|---|---|---|---|
| | **0.5** | **0.6** | **0.7** | **0.8** |

| | | | | |
|---|---|---|---|---|
| 1 | 0.95 | 0.31 | 0.00 | 0.00 |
| 2 | 0.70 | 0.06 | 0.00 | 0.00 |
| 3 | 0.95 | 0.38 | 0.01 | 0.00 |
| 4 | 0.82 | 0.36 | 0.06 | 0.00 |
| 5 | 0.90 | 0.46 | 0.07 | 0.00 |
| 6 | 0.95 | 0.61 | 0.03 | 0.00 |
| 7 | 0.97 | 0.29 | 0.00 | 0.00 |
| 8 | 0.96 | 0.47 | 0.02 | 0.00 |
| 9 | 0.91 | 0.32 | 0.01 | 0.00 |
| 10 | 0.90 | 0.32 | 0.01 | 0.00 |
| 11 | 0.99 | 0.56 | 0.09 | 0.00 |
| 12 | 0.93 | 0.62 | 0.06 | 0.00 |
| 13 | 1.00 | 0.96 | 0.53 | 0.02 |
| 14 | 0.94 | 0.86 | 0.41 | 0.04 |
| 15 | 0.84 | 0.27 | 0.02 | 0.00 |
| 16 | 0.99 | 0.90 | 0.46 | 0.02 |
| 17 | 1.00 | 0.98 | 0.60 | 0.06 |
| 18 | 0.94 | 0.59 | 0.14 | 0.01 |
| 19 | 0.91 | 0.65 | 0.09 | 0.01 |
| 20 | 0.92 | 0.60 | 0.10 | 0.01 |
| 21 | 0.96 | 0.48 | 0.00 | 0.00 |
| 22 | 0.89 | 0.56 | 0.10 | 0.00 |
| 23 | 0.98 | 0.50 | 0.04 | 0.00 |
| 24 | 0.99 | 0.62 | 0.00 | 0.00 |
| 25 | 1.00 | 0.89 | 0.24 | 0.00 |
| 26 | 0.96 | 0.66 | 0.19 | 0.01 |
| 27 | 0.99 | 0.90 | 0.12 | 0.00 |
| 28 | 0.98 | 0.66 | 0.02 | 0.00 |
| 29 | 0.98 | 0.77 | 0.17 | 0.01 |

| | | | | |
|---|---|---|---|---|
| 30 | 0.92 | 0.53 | 0.02 | 0.00 |

**Table S7.** Sequence novelty results of High-PepBinder

#### *8. Sequence novelty results of RFDiffusion*

| Target | Threshold | | | |
|---|---|---|---|---|
| | **0.5** | **0.6** | **0.7** | **0.8** |
| 1 | 0.91 | 0.27 | 0.02 | 0.01 |
| 2 | 0.62 | 0.17 | 0.06 | 0.00 |
| 3 | 0.84 | 0.30 | 0.11 | 0.00 |
| 4 | 0.93 | 0.46 | 0.15 | 0.02 |
| 5 | 0.92 | 0.47 | 0.09 | 0.01 |
| 6 | 0.98 | 0.81 | 0.03 | 0.00 |
| 7 | 0.80 | 0.05 | 0.00 | 0.00 |
| 8 | 0.95 | 0.39 | 0.00 | 0.00 |
| 9 | 0.99 | 0.95 | 0.62 | 0.30 |
| 10 | 0.77 | 0.18 | 0.00 | 0.00 |
| 11 | 1.00 | 0.94 | 0.58 | 0.10 |
| 12 | 0.94 | 0.75 | 0.00 | 0.00 |
| 13 | 1.00 | 0.75 | 0.16 | 0.00 |
| 14 | 0.98 | 0.84 | 0.25 | 0.05 |
| 15 | 0.96 | 0.65 | 0.03 | 0.01 |
| 16 | 1.00 | 0.91 | 0.10 | 0.00 |
| 17 | 1.00 | 1.00 | 0.83 | 0.20 |
| 18 | 1.00 | 0.91 | 0.53 | 0.03 |
| 19 | 0.95 | 0.73 | 0.11 | 0.00 |
| 20 | 0.99 | 0.75 | 0.28 | 0.00 |
| 21 | 0.95 | 0.34 | 0.00 | 0.00 |
| 22 | 0.97 | 0.67 | 0.09 | 0.00 |
| 23 | 0.89 | 0.22 | 0.02 | 0.00 |

| | | | | |
|---|---|---|---|---|
| 24 | 1.00 | 0.82 | 0.58 | 0.06 |
| 25 | 0.85 | 0.55 | 0.03 | 0.00 |
| 26 | 0.99 | 0.83 | 0.00 | 0.00 |
| 27 | 1.00 | 1.00 | 0.66 | 0.02 |
| 28 | 1.00 | 0.99 | 0.51 | 0.01 |
| 29 | 0.98 | 0.65 | 0.01 | 0.00 |
| 30 | 0.83 | 0.43 | 0.00 | 0.00 |

**Table S8.** Sequence novelty results of RFDiffusion

#### *9. Sequence novelty results of PepMLM*

| Target | Threshold | | | |
|---|---|---|---|---|
| | **0.5** | **0.6** | **0.7** | **0.8** |
| 1 | 0.56 | 0.12 | 0.00 | 0.00 |
| 2 | 0.67 | 0.19 | 0.00 | 0.00 |
| 3 | 0.34 | 0.00 | 0.00 | 0.00 |
| 4 | 0.28 | 0.06 | 0.00 | 0.00 |
| 5 | 0.54 | 0.02 | 0.00 | 0.00 |
| 6 | 0.22 | 0.00 | 0.00 | 0.00 |
| 7 | 0.46 | 0.00 | 0.00 | 0.00 |
| 8 | 0.61 | 0.03 | 0.00 | 0.00 |
| 9 | 0.88 | 0.13 | 0.00 | 0.00 |
| 10 | 0.88 | 0.12 | 0.00 | 0.00 |
| 11 | 0.99 | 0.86 | 0.19 | 0.00 |
| 12 | 0.70 | 0.20 | 0.00 | 0.00 |
| 13 | 1.00 | 0.85 | 0.21 | 0.00 |
| 14 | 0.41 | 0.00 | 0.00 | 0.00 |
| 15 | 0.94 | 0.51 | 0.01 | 0.00 |
| 16 | 0.37 | 0.07 | 0.00 | 0.00 |
| 17 | 1.00 | 0.99 | 0.26 | 0.00 |

| | | | | |
|---|---|---|---|---|
| 18 | 0.93 | 0.68 | 0.12 | 0.00 |
| 19 | 0.97 | 0.67 | 0.01 | 0.00 |
| 20 | 0.98 | 0.80 | 0.17 | 0.01 |
| 21 | 0.96 | 0.47 | 0.00 | 0.00 |
| 22 | 0.81 | 0.48 | 0.04 | 0.00 |
| 23 | 0.88 | 0.50 | 0.01 | 0.00 |
| 24 | 0.64 | 0.11 | 0.00 | 0.00 |
| 25 | 0.02 | 0.00 | 0.00 | 0.00 |
| 26 | 0.99 | 0.92 | 0.12 | 0.00 |
| 27 | 0.46 | 0.04 | 0.00 | 0.00 |
| 28 | 0.29 | 0.02 | 0.00 | 0.00 |
| 29 | 0.61 | 0.27 | 0.00 | 0.00 |
| 30 | 0.59 | 0.00 | 0.00 | 0.00 |

**Table S9.** Sequence novelty results of PepMLM

#### *10. Structure diversity results of High-PepBinder*

| **Target** | **Threshold** | | | |
|---|---|---|---|---|
| | **0.5** | **0.6** | **0.7** | **0.8** |
| 1 | 0.16 | 0.35 | 0.56 | 0.80 |
| 2 | 0.18 | 0.37 | 0.56 | 0.71 |
| 3 | 0.27 | 0.47 | 0.66 | 0.81 |
| 4 | 0.27 | 0.46 | 0.61 | 0.81 |
| 5 | 0.24 | 0.51 | 0.68 | 0.86 |
| 6 | 0.19 | 0.40 | 0.62 | 0.82 |
| 7 | 0.13 | 0.27 | 0.44 | 0.67 |
| 8 | 0.34 | 0.55 | 0.73 | 0.88 |
| 9 | 0.20 | 0.42 | 0.69 | 0.84 |
| 10 | 0.25 | 0.43 | 0.60 | 0.80 |
| 11 | 0.16 | 0.34 | 0.60 | 0.75 |

| | | | | |
|---|---|---|---|---|
| 12 | 0.60 | 0.76 | 0.88 | 0.95 |
| 13 | 0.18 | 0.43 | 0.64 | 0.81 |
| 14 | 0.47 | 0.71 | 0.85 | 0.94 |
| 15 | 0.39 | 0.54 | 0.68 | 0.84 |
| 16 | 0.32 | 0.52 | 0.72 | 0.87 |
| 17 | 0.19 | 0.45 | 0.65 | 0.85 |
| 18 | 0.24 | 0.42 | 0.63 | 0.83 |
| 19 | 0.31 | 0.52 | 0.68 | 0.77 |
| 20 | 0.57 | 0.73 | 0.86 | 0.94 |
| 21 | 0.44 | 0.62 | 0.75 | 0.90 |
| 22 | 0.34 | 0.53 | 0.69 | 0.86 |
| 23 | 0.43 | 0.64 | 0.75 | 0.89 |
| 24 | 0.32 | 0.59 | 0.79 | 0.89 |
| 25 | 0.39 | 0.60 | 0.80 | 0.89 |
| 26 | 0.13 | 0.40 | 0.62 | 0.80 |
| 27 | 0.39 | 0.61 | 0.76 | 0.90 |
| 28 | 0.69 | 0.84 | 0.96 | 0.99 |
| 29 | 0.42 | 0.56 | 0.74 | 0.89 |
| 30 | 0.46 | 0.70 | 0.90 | 0.96 |

**Table S10.** Structure diversity results of High-PepBinder

### *11. Structure diversity results of RFDiffusion*

| **Target** | **Threshold** | | | |
|---|---|---|---|---|
| | **0.5** | **0.6** | **0.7** | **0.8** |
| 1 | 0.02 | 0.06 | 0.19 | 0.43 |
| 2 | 0.01 | 0.02 | 0.06 | 0.18 |
| 3 | 0.43 | 0.62 | 0.76 | 0.86 |
| 4 | 0.29 | 0.51 | 0.63 | 0.66 |
| 5 | 0.41 | 0.54 | 0.66 | 0.81 |

| | | | | |
|---|---|---|---|---|
| 6 | 0.15 | 0.23 | 0.32 | 0.44 |
| 7 | 0.13 | 0.30 | 0.49 | 0.70 |
| 8 | 0.12 | 0.17 | 0.29 | 0.45 |
| 9 | 0.27 | 0.45 | 0.67 | 0.83 |
| 10 | 0.50 | 0.68 | 0.82 | 0.90 |
| 11 | 0.07 | 0.20 | 0.40 | 0.59 |
| 12 | 0.13 | 0.19 | 0.40 | 0.66 |
| 13 | 0.74 | 0.84 | 0.92 | 0.97 |
| 14 | 0.08 | 0.14 | 0.29 | 0.49 |
| 15 | 0.07 | 0.20 | 0.36 | 0.51 |
| 16 | 0.45 | 0.65 | 0.81 | 0.91 |
| 17 | 0.14 | 0.22 | 0.35 | 0.50 |
| 18 | 0.32 | 0.48 | 0.62 | 0.75 |
| 19 | 0.56 | 0.73 | 0.78 | 0.80 |
| 20 | 0.43 | 0.66 | 0.83 | 0.92 |
| 21 | 0.25 | 0.39 | 0.58 | 0.68 |
| 22 | 0.46 | 0.71 | 0.88 | 0.95 |
| 23 | 0.51 | 0.75 | 0.89 | 0.94 |
| 24 | 0.03 | 0.06 | 0.22 | 0.51 |
| 25 | 0.08 | 0.22 | 0.42 | 0.64 |
| 26 | 0.20 | 0.35 | 0.57 | 0.76 |
| 27 | 0.09 | 0.18 | 0.37 | 0.58 |
| 28 | 0.24 | 0.42 | 0.64 | 0.75 |
| 29 | 0.10 | 0.18 | 0.30 | 0.48 |
| 30 | 0.05 | 0.08 | 0.16 | 0.36 |

**Table S11.** Structure diversity results of RFDiffusion

#### *12. Structure diversity results of PepMLM*

| Target | Threshold |
|---|---|

| | **0.5** | **0.6** | **0.7** | **0.8** |
|---|---|---|---|---|
| 1 | 0.14 | 0.31 | 0.52 | 0.72 |
| 2 | 0.14 | 0.20 | 0.26 | 0.42 |
| 3 | 0.53 | 0.72 | 0.87 | 0.96 |
| 4 | 0.70 | 0.90 | 0.95 | 0.98 |
| 5 | 0.57 | 0.76 | 0.90 | 0.97 |
| 6 | 0.10 | 0.14 | 0.18 | 0.33 |
| 7 | 0.60 | 0.82 | 0.87 | 0.93 |
| 8 | 0.41 | 0.61 | 0.78 | 0.93 |
| 9 | 0.68 | 0.83 | 0.91 | 0.94 |
| 10 | 0.90 | 0.98 | 1.00 | 1.00 |
| 11 | 0.35 | 0.45 | 0.61 | 0.76 |
| 12 | 0.82 | 0.90 | 0.96 | 0.98 |
| 13 | 0.18 | 0.39 | 0.64 | 0.89 |
| 14 | 0.03 | 0.20 | 0.43 | 0.60 |
| 15 | 0.23 | 0.42 | 0.64 | 0.86 |
| 16 | 0.54 | 0.67 | 0.79 | 0.92 |
| 17 | 0.65 | 0.88 | 0.95 | 0.98 |
| 18 | 0.48 | 0.69 | 0.86 | 0.99 |
| 19 | 0.79 | 0.93 | 1.00 | 1.00 |
| 20 | 0.48 | 0.62 | 0.77 | 0.94 |
| 21 | 0.79 | 0.91 | 0.99 | 0.99 |
| 22 | 0.99 | 1.00 | 1.00 | 1.00 |
| 23 | 0.06 | 0.13 | 0.20 | 0.33 |
| 24 | 0.90 | 0.97 | 1.00 | 1.00 |
| 25 | 0.69 | 0.85 | 0.92 | 0.99 |
| 26 | 0.29 | 0.54 | 0.67 | 0.82 |
| 27 | 0.19 | 0.42 | 0.64 | 0.90 |
| 28 | 0.34 | 0.71 | 0.89 | 0.95 |

| | | | | |
|---|---|---|---|---|
| 29 | 0.34 | 0.52 | 0.63 | 0.81 |
| 30 | 0.18 | 0.41 | 0.57 | 0.83 |

**Table S12.** Structure diversity results of PepMLM

#### *13. Structure novelty results of High-PepBinder*

| Target | Threshold | | | |
|---|---|---|---|---|
| | **0.5** | **0.6** | **0.7** | **0.8** |
| 1 | 0.17 | 0.29 | 0.50 | 0.76 |
| 2 | 0.10 | 0.38 | 0.64 | 0.80 |
| 3 | 0.19 | 0.28 | 0.42 | 0.60 |
| 4 | 0.00 | 0.06 | 0.25 | 0.71 |
| 5 | 0.26 | 0.42 | 0.67 | 0.85 |
| 6 | 0.32 | 0.59 | 0.71 | 0.86 |
| 7 | 0.17 | 0.33 | 0.43 | 0.62 |
| 8 | 0.52 | 0.71 | 0.83 | 0.89 |
| 9 | 0.09 | 0.36 | 0.56 | 0.72 |
| 10 | 0.03 | 0.18 | 0.47 | 0.77 |
| 11 | 0.36 | 0.52 | 0.66 | 0.82 |
| 12 | 0.50 | 0.78 | 0.94 | 0.96 |
| 13 | 0.48 | 0.62 | 0.77 | 0.90 |
| 14 | 0.82 | 0.88 | 0.92 | 0.99 |
| 15 | 0.32 | 0.61 | 0.78 | 0.93 |
| 16 | 0.76 | 0.86 | 0.93 | 0.98 |
| 17 | 0.45 | 0.61 | 0.80 | 0.87 |
| 18 | 0.06 | 0.17 | 0.50 | 0.92 |
| 19 | 0.10 | 0.19 | 0.41 | 0.67 |
| 20 | 0.08 | 0.38 | 0.65 | 0.84 |
| 21 | 0.28 | 0.58 | 0.81 | 0.97 |
| 22 | 0.49 | 0.70 | 0.83 | 0.93 |

| | | | | |
|---|---|---|---|---|
| 23 | 0.02 | 0.07 | 0.55 | 0.87 |
| 24 | 0.65 | 0.78 | 0.88 | 0.93 |
| 25 | 0.59 | 0.72 | 0.88 | 0.95 |
| 26 | 0.26 | 0.40 | 0.61 | 0.76 |
| 27 | 0.82 | 0.93 | 0.96 | 1.00 |
| 28 | 0.89 | 0.97 | 0.99 | 0.99 |
| 29 | 0.01 | 0.26 | 0.64 | 0.91 |
| 30 | 0.34 | 0.56 | 0.79 | 0.96 |

**Table S13.** Structure novelty results of High-PepBinder

#### *14. Structure novelty results of RFDiffusion*

| **Target** | **Threshold** | | | |
|---|---|---|---|---|
| | **0.5** | **0.6** | **0.7** | **0.8** |
| 1 | 0.01 | 0.06 | 0.16 | 0.48 |
| 2 | 0.33 | 0.78 | 0.95 | 1.00 |
| 3 | 0.34 | 0.56 | 0.60 | 0.70 |
| 4 | 0.00 | 0.51 | 0.73 | 0.95 |
| 5 | 0.03 | 0.05 | 0.05 | 0.23 |
| 6 | 0.01 | 0.02 | 0.08 | 0.41 |
| 7 | 0.00 | 0.00 | 0.00 | 0.12 |
| 8 | 0.79 | 0.89 | 0.97 | 1.00 |
| 9 | 0.28 | 0.47 | 0.52 | 0.55 |
| 10 | 0.06 | 0.13 | 0.39 | 0.62 |
| 11 | 0.20 | 0.26 | 0.34 | 0.43 |
| 12 | 0.72 | 0.92 | 1.00 | 1.00 |
| 13 | 0.80 | 0.93 | 0.99 | 1.00 |
| 14 | 0.93 | 0.99 | 1.00 | 1.00 |
| 15 | 0.63 | 0.92 | 1.00 | 1.00 |
| 16 | 0.94 | 0.99 | 1.00 | 1.00 |

| | | | | |
|---|---|---|---|---|
| 17 | 0.92 | 0.95 | 0.99 | 1.00 |
| 18 | 0.00 | 0.05 | 0.23 | 0.82 |
| 19 | 0.45 | 0.62 | 0.70 | 0.82 |
| 20 | 0.08 | 0.44 | 0.79 | 0.92 |
| 21 | 0.05 | 0.24 | 0.64 | 0.98 |
| 22 | 0.70 | 1.00 | 1.00 | 1.00 |
| 23 | 0.03 | 0.10 | 0.55 | 0.92 |
| 24 | 0.89 | 0.99 | 0.99 | 1.00 |
| 25 | 0.90 | 0.95 | 1.00 | 1.00 |
| 26 | 0.46 | 0.63 | 0.73 | 0.81 |
| 27 | 0.87 | 0.98 | 1.00 | 1.00 |
| 28 | 0.96 | 1.00 | 1.00 | 1.00 |
| 29 | 0.00 | 0.06 | 0.55 | 0.97 |
| 30 | 0.04 | 0.06 | 0.59 | 0.93 |

**Table S14.** Structure novelty results of RFDiffusion

#### *15. Structure novelty results of PepMLM*

| **Target** | **Threshold** | | | |
|---|---|---|---|---|
| | **0.5** | **0.6** | **0.7** | **0.8** |
| 1 | 0.23 | 0.29 | 0.36 | 0.54 |
| 2 | 0.51 | 0.89 | 0.96 | 1.00 |
| 3 | 0.01 | 0.04 | 0.12 | 0.21 |
| 4 | 0.00 | 0.10 | 0.16 | 0.55 |
| 5 | 0.01 | 0.01 | 0.09 | 0.47 |
| 6 | 0.01 | 0.04 | 0.12 | 0.36 |
| 7 | 0.09 | 0.17 | 0.21 | 0.35 |
| 8 | 0.66 | 0.81 | 0.95 | 0.98 |
| 9 | 0.11 | 0.47 | 0.64 | 0.82 |
| 10 | 0.02 | 0.08 | 0.23 | 0.60 |

| | | | | |
|---|---|---|---|---|
| 11 | 0.07 | 0.11 | 0.15 | 0.37 |
| 12 | 0.48 | 0.87 | 0.95 | 0.99 |
| 13 | 0.74 | 0.79 | 0.85 | 0.99 |
| 14 | 0.90 | 0.95 | 0.99 | 1.00 |
| 15 | 0.81 | 0.97 | 0.99 | 1.00 |
| 16 | 0.88 | 0.96 | 0.99 | 0.99 |
| 17 | 0.79 | 0.85 | 0.94 | 0.99 |
| 18 | 0.00 | 0.03 | 0.28 | 0.87 |
| 19 | 0.33 | 0.49 | 0.66 | 0.85 |
| 20 | 0.18 | 0.53 | 0.76 | 0.96 |
| 21 | 0.21 | 0.61 | 0.89 | 1.00 |
| 22 | 0.72 | 0.94 | 0.99 | 1.00 |
| 23 | 0.03 | 0.07 | 0.45 | 0.82 |
| 24 | 0.95 | 0.97 | 1.00 | 1.00 |
| 25 | 1.00 | 1.00 | 1.00 | 1.00 |
| 26 | 0.01 | 0.02 | 0.02 | 0.04 |
| 27 | 0.93 | 0.97 | 0.98 | 0.99 |
| 28 | 0.81 | 0.88 | 0.95 | 1.00 |
| 29 | 0.00 | 0.06 | 0.21 | 0.45 |
| 30 | 0.04 | 0.12 | 0.55 | 0.90 |

**Table S15.** Structure novelty results of PepMLM

### Section S6. Results on the Binding Capability of Generated Peptides to Target Proteins

#### *1. Mean ipTM*

| **Target** | **High-PepBinder** | **RFDiffusion** | **PepMLM** |
|---|---|---|---|
| 1 | 0.634 | 0.609 | 0.794 |

| | | | |
|---|---|---|---|
| 2 | 0.532 | 0.598 | 0.483 |
| 3 | 0.581 | 0.454 | 0.587 |
| 4 | 0.402 | 0.256 | 0.376 |
| 5 | 0.636 | 0.744 | 0.735 |
| 6 | 0.350 | 0.327 | 0.361 |
| 7 | 0.569 | 0.560 | 0.561 |
| 8 | 0.603 | 0.553 | 0.513 |
| 9 | 0.670 | 0.593 | 0.522 |
| 10 | 0.714 | 0.500 | 0.479 |
| 11 | 0.527 | 0.485 | 0.501 |
| 12 | 0.613 | 0.524 | 0.466 |
| 13 | 0.517 | 0.316 | 0.334 |
| 14 | 0.265 | 0.421 | 0.320 |
| 15 | 0.635 | 0.645 | 0.495 |
| 16 | 0.376 | 0.322 | 0.381 |
| 17 | 0.404 | 0.360 | 0.356 |
| 18 | 0.595 | 0.519 | 0.572 |
| 19 | 0.789 | 0.638 | 0.618 |
| 20 | 0.626 | 0.681 | 0.566 |
| 21 | 0.547 | 0.381 | 0.415 |
| 22 | 0.604 | 0.569 | 0.496 |
| 23 | 0.610 | 0.610 | 0.509 |
| 24 | 0.548 | 0.650 | 0.582 |
| 25 | 0.512 | 0.518 | 0.438 |
| 26 | 0.543 | 0.630 | 0.669 |
| 27 | 0.490 | 0.601 | 0.570 |
| 28 | 0.565 | 0.480 | 0.719 |
| 29 | 0.677 | 0.498 | 0.621 |
| 30 | 0.596 | 0.624 | 0.518 |

**Table S16.** Mean ipTM results

#### *2. Max ipTM*

| Target | High-PepBinder | RFDiffusion | PepMLM |
|---|---|---|---|
| 1 | 0.82 | 0.86 | 0.92 |
| 2 | 0.77 | 0.88 | 0.81 |
| 3 | 0.94 | 0.85 | 0.92 |
| 4 | 0.71 | 0.63 | 0.79 |
| 5 | 0.94 | 0.93 | 0.87 |
| 6 | 0.65 | 0.86 | 0.57 |
| 7 | 0.83 | 0.82 | 0.83 |
| 8 | 0.92 | 0.89 | 0.8 |
| 9 | 0.82 | 0.81 | 0.83 |
| 10 | 0.96 | 0.95 | 0.89 |
| 11 | 0.93 | 0.93 | 0.95 |
| 12 | 0.84 | 0.77 | 0.73 |
| 13 | 0.86 | 0.61 | 0.81 |
| 14 | 0.65 | 0.75 | 0.59 |
| 15 | 0.95 | 0.9 | 0.89 |
| 16 | 0.83 | 0.55 | 0.75 |
| 17 | 0.87 | 0.81 | 0.74 |
| 18 | 0.94 | 0.87 | 0.87 |
| 19 | 0.96 | 0.95 | 0.92 |
| 20 | 0.85 | 0.88 | 0.82 |
| 21 | 0.81 | 0.71 | 0.72 |
| 22 | 0.84 | 0.87 | 0.68 |
| 23 | 0.97 | 0.96 | 0.96 |
| 24 | 0.78 | 0.89 | 0.87 |

| | | | |
|---|---|---|---|
| 25 | 0.72 | 0.73 | 0.64 |
| 26 | 0.90 | 0.89 | 0.91 |
| 27 | 0.83 | 0.89 | 0.91 |
| 28 | 0.84 | 0.77 | 0.9 |
| 29 | 0.93 | 0.92 | 0.96 |
| 30 | 0.87 | 0.87 | 0.88 |

**Table S17.** Max ipTM results

#### *3. Median ipAE*

| Target | High-PepBinder | RFDiffusion | PepMLM |
|---|---|---|---|
| 1 | 5.178 | 6.415 | 3.399 |
| 2 | 9.167 | 8.093 | 10.151 |
| 3 | 10.897 | 14.420 | 10.808 |
| 4 | 13.879 | 18.270 | 14.906 |
| 5 | 6.177 | 4.326 | 4.609 |
| 6 | 15.854 | 16.266 | 14.576 |
| 7 | 7.706 | 6.958 | 8.296 |
| 8 | 14.884 | 15.138 | 17.639 |
| 9 | 3.559 | 4.478 | 6.232 |
| 10 | 7.359 | 12.979 | 13.523 |
| 11 | 11.190 | 12.260 | 12.455 |
| 12 | 5.822 | 7.407 | 9.505 |
| 13 | 15.210 | 21.265 | 20.708 |
| 14 | 10.702 | 7.260 | 9.607 |
| 15 | 11.638 | 11.539 | 16.384 |
| 16 | 19.553 | 20.799 | 18.984 |
| 17 | 18.687 | 19.880 | 19.968 |
| 18 | 7.764 | 8.581 | 8.138 |

| | | | |
|---|---|---|---|
| 19 | 5.480 | 9.393 | 9.693 |
| 20 | 7.128 | 5.462 | 6.689 |
| 21 | 14.389 | 18.564 | 18.508 |
| 22 | 7.423 | 8.165 | 9.280 |
| 23 | 12.913 | 14.098 | 16.590 |
| 24 | 6.797 | 4.884 | 6.001 |
| 25 | 7.215 | 6.788 | 8.782 |
| 26 | 7.803 | 6.650 | 5.412 |
| 27 | 9.119 | 7.006 | 7.454 |
| 28 | 12.137 | 14.930 | 9.368 |
| 29 | 8.723 | 13.145 | 10.202 |
| 30 | 7.078 | 7.086 | 8.967 |

**Table S18.** Median ipAE results

#### *4. Hit Rate*

We define a hit as any generated peptide whose ipTM value exceeds that of the reference peptide. The hit rate is calculated as the number of hits divided by the total number of generated peptides.

| Target | High-PepBinder | RFDiffusion | PepMLM |
|---|---|---|---|
| 1 | 0.00 | 0.00 | 0.14 |
| 2 | 0.33 | 0.49 | 0.17 |
| 3 | 0.10 | 0.05 | 0.13 |
| 4 | 0.10 | 0.01 | 0.02 |
| 5 | 0.04 | 0.10 | 0.00 |
| 6 | 0.69 | 0.60 | 0.68 |
| 7 | 0.12 | 0.21 | 0.14 |
| 8 | 0.56 | 0.42 | 0.37 |

| | | | |
|---|---|---|---|
| 9 | 0.07 | 0.06 | 0.02 |
| 10 | 0.05 | 0.02 | 0.00 |
| 11 | 0.04 | 0.03 | 0.06 |
| 12 | 0.02 | 0.01 | 0.00 |
| 13 | 0.82 | 0.54 | 0.54 |
| 14 | 0.38 | 0.79 | 0.55 |
| 15 | 0.04 | 0.00 | 0.00 |
| 16 | 0.16 | 0.02 | 0.11 |
| 17 | 0.53 | 0.41 | 0.29 |
| 18 | 0.31 | 0.06 | 0.18 |
| 19 | 0.15 | 0.09 | 0.01 |
| 20 | 0.15 | 0.21 | 0.07 |
| 21 | 0.23 | 0.04 | 0.04 |
| 22 | 0.70 | 0.48 | 0.21 |
| 23 | 0.02 | 0.02 | 0.02 |
| 24 | 0.09 | 0.39 | 0.17 |
| 25 | 0.10 | 0.05 | 0.08 |
| 26 | 0.00 | 0.00 | 0.00 |
| 27 | 0.02 | 0.13 | 0.08 |
| 28 | 0.15 | 0.03 | 0.57 |
| 29 | 0.02 | 0.01 | 0.12 |
| 30 | 0.00 | 0.00 | 0.00 |

**Table S19.** Hit Rate results

#### *5. The proportion of samples with Rosetta dG_separated/dSASA ×100 < -1.5*

| Target | High-PepBinder | RFDiffusion | PepMLM |
|---|---|---|---|
| 1 | 0.41 | 0.51 | 0.80 |
| 2 | 0.25 | 0.67 | 0.27 |

| | | | |
|---|---|---|---|
| 3 | 0.50 | 0.45 | 0.47 |
| 4 | 0.22 | 0.34 | 0.11 |
| 5 | 0.83 | 0.82 | 0.87 |
| 6 | 0.59 | 0.50 | 0.43 |
| 7 | 0.16 | 0.17 | 0.21 |
| 8 | 0.22 | 0.43 | 0.04 |
| 9 | 0.12 | 0.05 | 0.09 |
| 10 | 0.39 | 0.34 | 0.18 |
| 11 | 0.43 | 0.34 | 0.22 |
| 12 | 0.03 | 0.02 | 0.02 |
| 13 | 0.15 | 0.12 | 0.07 |
| 14 | 0.10 | 0.12 | 0.06 |
| 15 | 0.35 | 0.23 | 0.12 |
| 16 | 0.10 | 0.05 | 0.06 |
| 17 | 0.06 | 0.19 | 0.03 |
| 18 | 0.40 | 0.27 | 0.39 |
| 19 | 0.52 | 0.48 | 0.53 |
| 20 | 0.42 | 0.34 | 0.49 |
| 21 | 0.23 | 0.29 | 0.25 |
| 22 | 0.07 | 0.02 | 0.01 |
| 23 | 0.23 | 0.22 | 0.11 |
| 24 | 0.18 | 0.22 | 0.17 |
| 25 | 0.04 | 0.01 | 0.00 |
| 26 | 0.29 | 0.52 | 0.45 |
| 27 | 0.25 | 0.30 | 0.36 |
| 28 | 0.57 | 0.26 | 0.47 |
| 29 | 0.42 | 0.25 | 0.61 |
| 30 | 0.42 | 0.37 | 0.36 |

**Table S20.** The proportion of samples with Rosetta dG_separated/dSASA $\times$ 100 < -1.5

### S7. Design of Peptide Binders Targeting TRAF6

TRAF6 is an intracellular signal transduction protein. Structure 1LB5 represents the reference complex between TRAF6 and the intracellular domain of the RANK receptor. Binders 1-3 are candidate peptides generated by High-PepBinder.

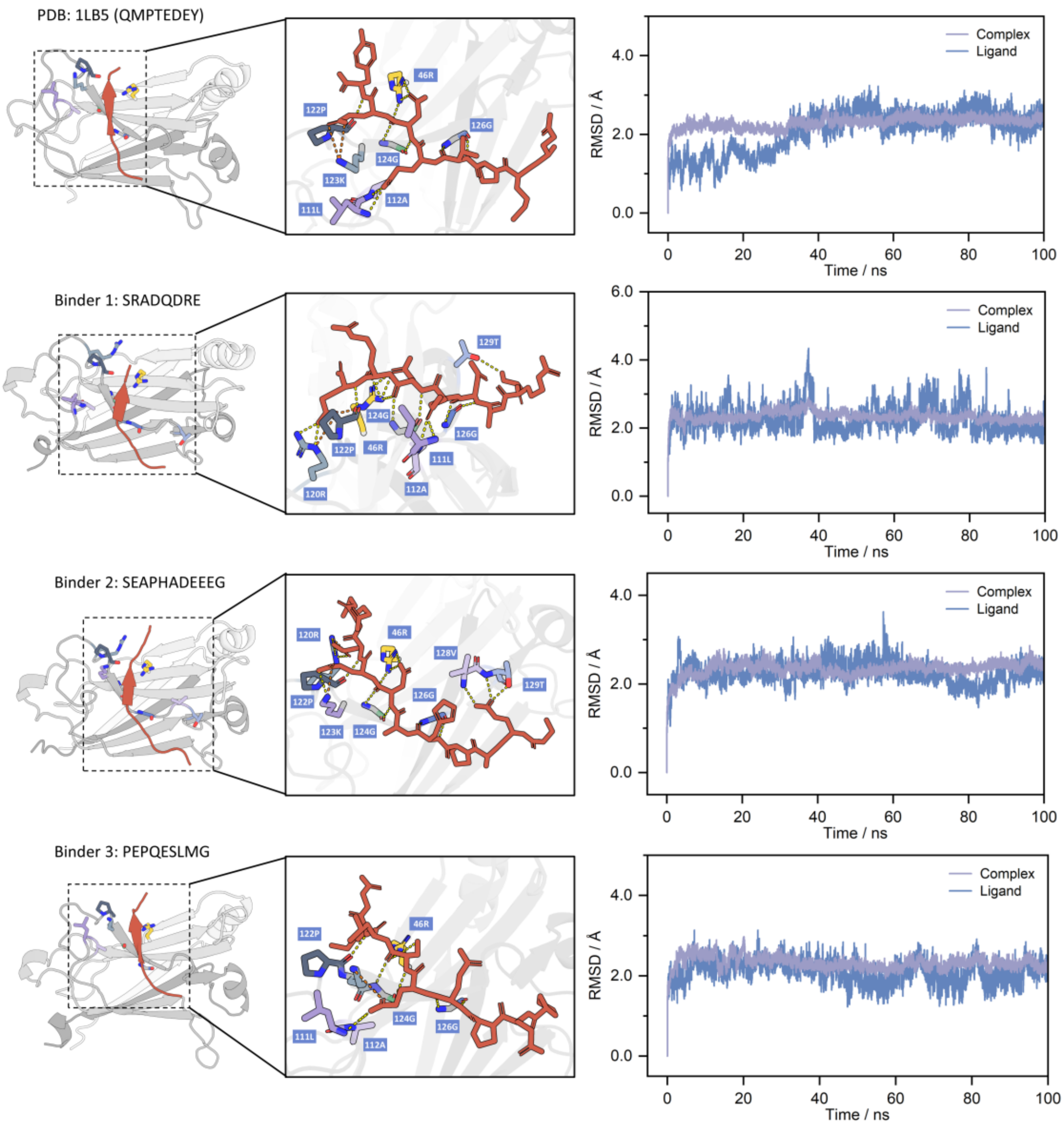

**Figure S1.** Peptide generation results for the TRAF6 target. The left panel shows the overall complex structure and schematic illustrations of secondary interactions at the binding site, where yellow indicates hydrogen bonds and orange indicates salt bridges. The right panel presents the root-mean-square deviation (RMSD) of the complex and the ligand during molecular dynamics simulations, which is used to assess binding stability.

| | ipTM | ipLDDT | ipAE | dG_separated/dSASA×100 (Å²) | H Bond | Salt Bridge | $\Delta G_{bind}$ (kcal/mol) |
|---|---|---|---|---|---|---|---|
| **Binder 1** | 0.94 | 87.70 | 1.7 | -1.703 | 13 | 2 | -65.04±3.57 |
| **Binder 2** | 0.89 | 85.86 | 2.2 | -2.007 | 15 | 2 | -67.09±4.01 |
| **Binder 3** | 0.94 | 91.96 | 1.6 | -1.678 | 9 | 1 | -67.76±3.98 |

**Table S21**. Results of three candidate peptides targeting TRAF6, where the reference complex (PDB: 1LB5) has a $\Delta G_{bind}$ of -62.96±3.57 kcal/mol and forms 10 hydrogen bonds and 2 salt bridges.

### S8. Design of Peptide Binders Targeting ESRRA

ESRRA is a nuclear receptor whose ligand binding typically relies on peptides containing a specific consensus motif, namely the canonical LXXLL motif, where X denotes any amino acid. This motif is a key determinant for the recognition and binding of nuclear receptor co-regulators. Structure 1XB7 represents the reference complex between the ligand-binding domain of ESRRA and a PGC-1α peptide. Binders 1-3 are candidate peptides generated by High-PepBinder, all of which contain the LXXLL motif.

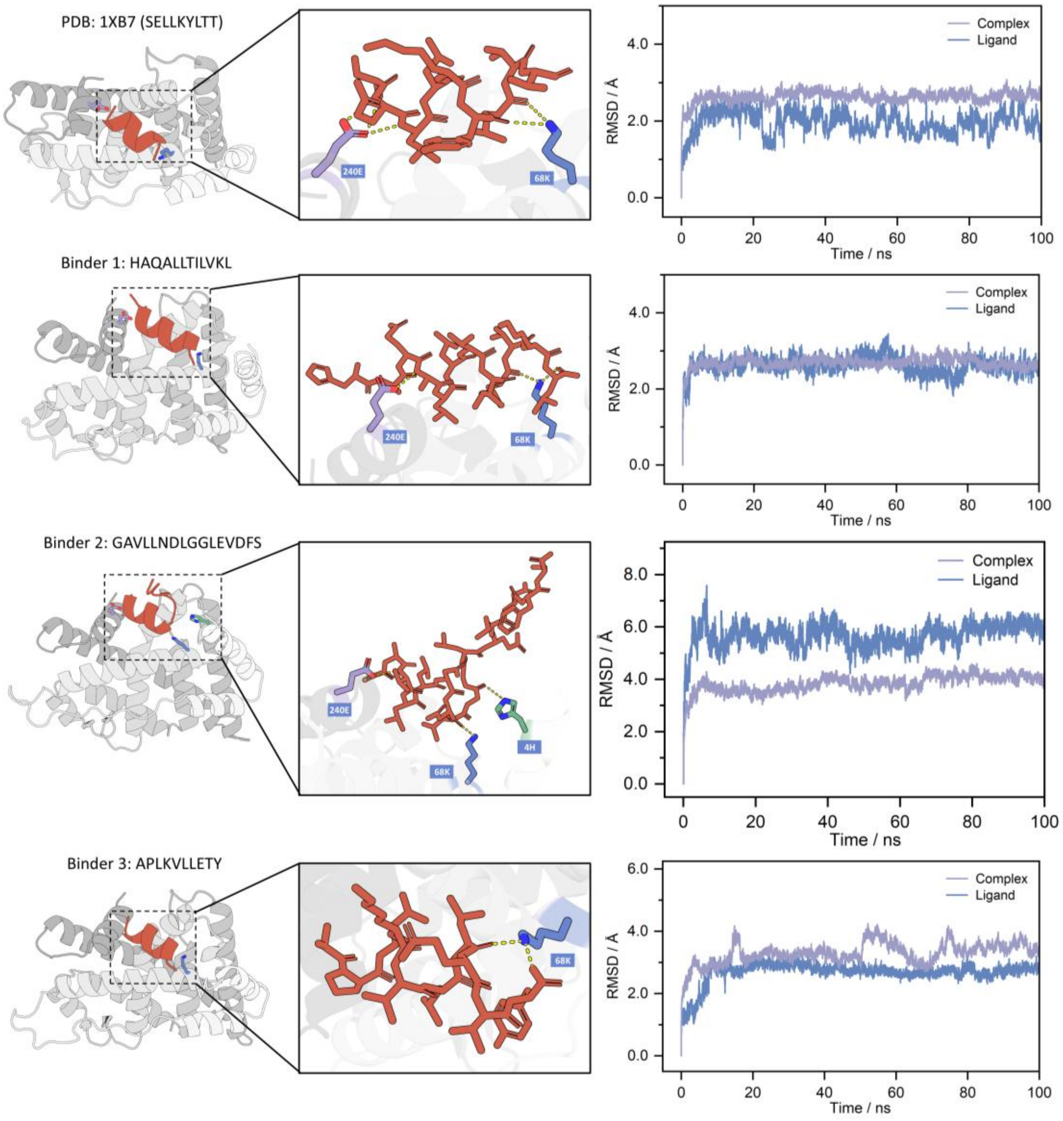

**Figure S2.** Peptide generation results for the ESRRA target.

| | ipTM | ipLDDT | ipAE | dG_separated/dSASA×100 (Å²) | H Bond | $\Delta G_{bind}$ (kcal/mol) |
|---|---|---|---|---|---|---|
| **Binder 1** | 0.91 | 86.51 | 2.3 | -2.520 | 3 | -81.90±5.09 |
| **Binder 2** | 0.81 | 74.83 | 2.5 | -2.901 | 4 | -81.18±4.89 |
| **Binder 3** | 0.93 | 91.24 | 2.0 | -2.526 | 2 | -54.72±4.72 |

**Table S22**. Results of three candidate peptides targeting ESRRA, where the reference complex (PDB: 1XB7) has a $\Delta G_{bind}$ of -54.36±5.36 kcal/mol and forms 5 hydrogen bonds.
